# Using induced pluripotent stem cells to investigate human neuronal phenotypes in 1q21.1 deletion and duplication syndrome

**DOI:** 10.1101/2021.02.08.430246

**Authors:** Gareth Chapman, Mouhamed Alsaqati, Sharna Lunn, Tanya Singh, Stefanie C Linden, David E. J. Linden, Marianne B.M. van den Bree, Mike Ziller, Michael J Owen, Jeremy Hall, Adrian J. Harwood, Yasir Ahmed Syed

**Affiliations:** Neuroscience and Mental Health Research Institute, Hadyn Ellis Building, Cathays, Cardiff, CF24 4HQ; School of Bioscience, The Sir Martin Evans Building, Museum Ave, Cardiff CF10 3AX; Division of Psychological Medicine and Clinical Neurosciences (DPMCN), School of Medicine, Cardiff University, Cardiff, UK; School of Mental Health and Neuroscience, Faculty of Health, Medicine and Life Sciences, Maastricht University, Maastricht, The Netherlands; MaxPlanck Institute for Psychiatry, Munich, Germany

## Abstract

Copy Number Variation (CNV) at the 1q21.1 locus is associated with a range of neurodevelopmental and psychiatric disorders in humans, including abnormalities in head size and motor deficits. Yet, the functional consequences of these CNVs (both deletion and duplication) on neuronal development remain unknown. To determine the impact of CNV at the 1q21.1 locus on neuronal development, we generated induced pluripotent stem cells from individuals harbouring 1q21.1 deletion or duplication and differentiated them into functional cortical neurons. We show that neurons with 1q21.1 deletion or duplication display reciprocal phenotype with respect to proliferation, differentiation potential, neuronal maturation, synaptic density, and functional activity. Deletion of the 1q21.1 locus was also associated with an increased expression of lower cortical layer markers. This difference was conserved in the mouse model of 1q21.1 deletion, which displayed altered corticogenesis. Importantly, we show that neurons with 1q21.1 deletion and duplication are associated with differential expression of calcium channels and demonstrate that physiological deficits in neurons with 1q21.1 deletion or duplication can be pharmacologically modulated by targeting Ca^2+^ channel activity. These findings provide biological insight into the neuropathological mechanism underlying 1q21.1 associated brain disorder and indicate a potential target for therapeutic interventions.

## Introduction

Investigating the biology of rare but relatively penetrant copy number variants (CNVs), provides an opportunity to understand the genetic basis of an increased susceptibility to a range of neurodevelopmental and neuropsychiatric disorders such as schizophrenia, autism, mental retardation and epilepsy^1-7^. There are now several prominent examples of pathogenic CNVs such as 1q21.1 deletions and duplications; 3q29 microduplications; 15q13.3 deletions; 16p11.2 deletions and duplications, and 22q11.2 deletions all of which are associated with increased risk for neurodevelopmental and neuropsychiatric disorders^8-10^. These CNVs are variable in size and can be either *de novo* or familial^11, 12^. Furthermore, a recent study showed that the brain is the tissue which is most intolerant to CNV associated changes in gene dosage^13^. Therefore, studying the impact of these CNVs on brain development provides a window of opportunity to understand the cellular mechanisms underlying increased risk for psychiatric disorders.

The 1q21.1 chromosomal locus (chr1: 146.57-147.39; GRCh37/hg19) contains at least four low copy repeats which render this region susceptible to non-allelic homologous recombination leading to recurrent deletions and duplications^14, 15,16^. Although its prevalence worldwide is not clear, data from UK Biobank has provided estimates of a population frequency of 0.027% for the 1q21.1 deletion and 0.044% for 1q21.1 duplication^17^. Two main classes of the 1q21.1 CNVs has been described. The more common Class I comprises the critical or distal region, whereas Class II compromises of the Thrombocytopenia Absent Radius (TAR) region in addition to the critical region^15, 18^. The critical/distal region is approximately 1.36Mb (from 145 to 146.35 Mb, according to NCBI build 36) and contain at least 12 protein coding genes including *PRKAB2, CHD1L, BCL9, ACP6, GJA5, GJA8 and NOTCH2NL*^9^. Phenotypes associated with distal 1q21.1 deletion include developmental delay, cognitive impairment, microcephaly, facial anomalies, schizophrenia, attention deficit hyperactivity disorder, emotional and behavioural problems. Whereas 1q21.1 distal duplication has been associated with macrocephaly, developmental delay, autism spectrum disorder, cognitive impairment, hypertelorism, and congenital cardiac anomalies^14, 15, 19-21^. Therefore, variation at this locus represents a clear risk factor for a range of neuropsychiatric disorders and need to be functionally characterised to understand the contribution of this loci to neurodevelopmental deficits leading to associated developmental psychiatric disorders. So far, the contribution of concomitantly deleted or duplicated genes in this locus towards the pathogenies of neuropsychiatric disorders is largely unknown.

To understand the impact of the Class I 1q21.1 CNV (from here referred to as 1q21.1 deletions or duplications) on neuronal development, we established a cellular model of by deriving human induced pluripotent stem cells (iPSCs) from subjects carrying 1q21.1 deletion or duplication and differentiated them into cortical neurons. We demonstrate that neural progenitor cells (NPCs) carrying 1q21.1 deletion or duplication are associated with early neurodevelopmental phenotypes. Furthermore, these NPCs after differentiation into neurons show dysregulated neuronal development, associated with altered morphology and synaptic density in comparison to controls. Moreover, these neurons are associated with dysregulated cortical layer identity. We validated aspects of these cellular phenotypes in a 1q21.1 microdeletion mouse model and show that some of these differences are conserved across species. Furthermore, we demonstrate that the presence of 1q21.1 CNVs impact the physiological and electrical properties of neurons as measured by calcium activity and multi-electrode arrays (MEAs). Finally, using iPSC derived neurons with 1q21.1 CNVs as an *in vitro* pharmacological model, we show that the aberrant physiological activity of these cells can be modulated by targeting Ca^2+^ channels.

## Methods

### Ethics statement

Generation and use of human iPSC were approved by the Cardiff University and HSE (GMO130/19.3). Clinical and psychometric testing (supplementary table 1) of participants and skin biopsies was approved by the Regional Ethics Committee of the National Health Service (study 14/WA/0035).

### iPSC generation, characterization and maintenance

Fibroblasts with subject carrying 1q21.1 deletion(n=3) or duplication (n=2) were reprogrammed into induced pluripotent stem cells (iPSCs) using the CytoTune™-IPS 2.0 Sendai reprogramming kit (Thermo-Fisher). Two established iPSC lines were used as controls (IBJ4 see Plumbly *et al*^*22*^ and HPSI1013i-wuye_2 purchased from HipSci). Pluripotency was confirmed by immunofluorescence, qPCR and trilineage differentiation (Supplementary Fig 1-5). iPSCs were grown on Geltrex™ coated plates in Essential 8™ Flex media. The cell lines were genotyped to identify the location of 1q21.1 locus and to identify any pathogenetic CNVs. Further, the cell lines were regularly tested to check any mycoplasma contamination.

### Cortical Neuronal differentiation

iPSCs were differentiated into cortical neurons using a modified version of a previously described protocol^23^. Cells were maintained until 90-100% confluent at which point the media was changed to N2B27-(2/3 DMEM/F12, 1/3 Neurobasal, N2 supplement, B27 supplement without retinoic acid, penicillin, streptomycin, glutamine and β-mercaptoethanol) supplemented with 250nM LDN-193189 (LDN) and 10µM SB431542 (SB). For the subsequent 10 days cells were maintained with both SB and LDN and then they were passaged onto fibronectin. Cells were maintained on fibronectin for 10 days in un-supplemented N2B27-media with ½ media changes every other day. Cells were then plated onto laminin and poly-D-lysine coated plates and after 2 days the media was replaced with N2B27+ (2/3 DMEM/F12, 1/3 Neurobasal, N2 supplement, B27 supplement, penicillin, streptomycin, glutamine and β-mercaptoethanol) after a further 2 days media was replaced with N2B27+ supplemented with CultureOne™ supplement. After 2 days media was replaced with fresh N2B27^+^ supplemented with 5µM DAPT and 1µM PD0332991 (PD). Cells were maintained with DAPT and PD for 4 days. Cells were then dissociated using Accutase® Solution and were re-plated on laminin and poly-D-lysine coated plates at a density of 200,000 cells/cm^2^. Cells were maintained in un-supplemented N2B27+ for up to 20 days with ½ media changes performed every other day. A minimum of three independent neuronal differentiation of all iPSC lines were done for the all the experiments reported.

### Calcium Imaging

Day 40 neuronal differentiations were passaged onto poly-d-lysine and laminin coated coverslips at a density of 50,000 cells/cm^2^. For pharmacological interventions, drugs (50nM verapamil and 2.5µM roscovitine) were applied 1 day after passaging and concentrations were maintained during all media changes (every 3 days). After 6 days media was replaced with BPM (BrainPhys™ Neuronal Medium supplemented with B27+ supplement, penicillin, streptomycin, glutamine, 10ng/mL BDNF and 35 µg/mL ascorbic acid). After 50 days of differentiation neurons were loaded with 1µM Cal-520® (AAT Bioquest) for 1 hour in BPM containing 0.02% Pluronic F127. The media was then replaced with fresh BPM and cells were incubated for a further 1 hour at 37°C. Coverslips were transferred into artificial cerebrospinal fluid (aSCF) containing: 125mM NaCl, 26mM NaHCO_3_, 1.25mM KH_2_PO_4_, 2.5mM KCl, 1mM MgCl_2_, 2mM CaCl_2_ and 25mM Glucose with or without either DL-AP5 or NQBX (at a final concentration of 10µM). Images were taken using an epifluorescence microscope at intervals of 200ms for 5 minutes. A minimum of 3 technical replicates (separate image stacks from a single culture) were averaged to generate each data point. Images were analysed using NeuroCa and Matlab. After automated analysis events with a rise time or fall time of more than 2 seconds were discarded and for the purpose of analysing average number of events all cells with no events were also discarded.

### Multiple Electrode Arrays

All experiments were performed using CytoView MEA 24-well plates (M384-tMEA-24W). MEAs were first pre-treated with 0.01% polyethylenimine (Sigma) and incubated for 1 hour at 37°C. Day 50 neurons were plated as high density drop cultures (5,000 cells/µL) containing 10µg/ml laminin. After 1-hour, conditioned medium was added into each MEA and after 24 hours 0.5ml of fresh BPM was added to each array. MEA cultures were maintained in 1:1 fresh to astrocyte condition BPM replaced every 3-4 days. Electrophysiological activity was recorded every 10 days using hardware (Maestro Pro complete with Maestro 768-channel amplifier) and software (AxIS 1.5.2) from Axion Biosystems (Axion Biosystems Inc., Atlanta, GA). Channels were sampled simultaneously with a gain of 1000× and a sampling rate of 12.5 kHz/channel. During the recording, the temperature was maintained constant at 37°C. A Butterworth band-pass filter (with a high-pass cut-off of 200 Hz and low-pass cut-off of 3000Hz) was applied along with a variable threshold spike detector set at 5.5× standard deviation on each channel. Offline analysis was achieved with custom scripts written in MATLAB (available on request). Briefly, spikes were detected from filtered data using an automatic threshold-based method set at -5.5 x σ, where σ is an estimate of the noise of each electrode based upon the median absolute deviation 1. Spike timestamps were analysed to provide statistics on the general excitability of cultures. Neuronal bursting was detected based on three parameters: inter-burst period longer than 200ms, more than 3 spikes in each burst and a maximum inter-spike (intra-burst) interval of 300 milliseconds. Network activity was illustrated by creating array–wide spike detection rate (ASDR) plots with a bin width of 200 ms. Synchronised bursts (SBs) across all electrodes in the network were identified using Axion built-in neural metric analysis tool employing the envelope algorithm. The algorithm defines a SB by identifying times when the histogram exceeds a threshold of 1 standard deviation above or below the mean with a minimum of 200ms between SBs and at least 10% of electrodes included. All active electrodes were included in the analysis. A minimum of three MEAs/cell line/differentiation have been considered for the analysis.

### Gene expression analysis

Primers for all target genes are listed in Supplemental Table 2. See Supplementary Methods for details.

### Western blotting

See Supplementary Methods for details

### Immunofluorescence and cell counting

See Supplementary Methods for details

### Histological analysis of mice brains

See Supplementary Methods for details

### Statistical analyses

Data is expressed as mean±SEM. All data is comprised of a minimum of 3 separate differentiations (n) for each cell line used in this study (2 control, 3 1q21.1 deletion and 2 1q21.1 duplication). All technical replicates were averaged before statistical testing. Statistical analyses were conducted in GraphPad Prism 6.01 (GraphPad Software). Differences between conditions or groups were evaluated using two-tailed unpaired Students T-Test or one/two-way ANOVA. *p*-values <0.05 were considered statistically significant.

## Results

### Deletions and duplications of the 1q21.1 locus is associated with altered neuronal development

We first assessed the effect of 1q21.1 deletion or duplication on the expression of genes within the distal 1q21.1 region, focusing on the expression of five key genes within this locus. After 50 days of differentiation three of these critical genes (BCL9, CDH1L and PRKAB2) had altered expression in 1q21.1 deletion or duplication in comparison to controls (Supplementary Fig 6B). To determine if deletion or duplication of the 1q21.1 locus altered neurodevelopmental trajectories we quantified the expression level of: a neural stem cell marker (NESTIN^24^); a marker of immature neurons Doublecortin (DCX^25^) and a mature neuronal marker (MAP2^26^) throughout the course of neuronal differentiation. The overall expression profile of NESTIN, Doublecortin (DCX) and MAP2 in controls was in accordance with previously published studies^27, 28^. NESTIN expression was significantly higher in 1q21.1 duplication, but unchanged in the 1q21.1 deletion group after 20 days of differentiation (Fig. 1B). Similarly, a second NPC marker, PLZF, was also elevated in duplications, but no change was seen for a third marker ZO1 in either duplications or deletions (Supplementary Fig 7B). However, these differences in mRNA did not translate into altered NESTIN+ cells at day 20(Fig. 1M). To examine the impact of 1q21.1 CNVs on proliferation potential of cells, we examined the expression of the mitotic marker Ki67. A small, but significant decrease of Ki67+ cells was observed in 1q21.1 deletion culture at day 20 and a substantial decrease in Ki67 mRNA was seen at day 30 for 1q21.1 deletion group (Fig 1 F-I, Supplementary Fig 7D). At day 30 of development, 1q21.1 duplication culture exhibit at reciprocal expression pattern of elevated Ki67 mRNA (Supplementary Fig 7D). Further, 1q21.1 deletion group expressed significantly lower levels of NOTCH2NL to that of control and duplication cells at day 20 of differentiation. Although this difference was also observed at day 50, it did not reach statistical significance (Supplementary Fig 6 B, C), suggesting that NOTCH2NL expression could be associated with early neurodevelopmental phenotypes associated with 1q21.1 CNV.

**Fig 1.**
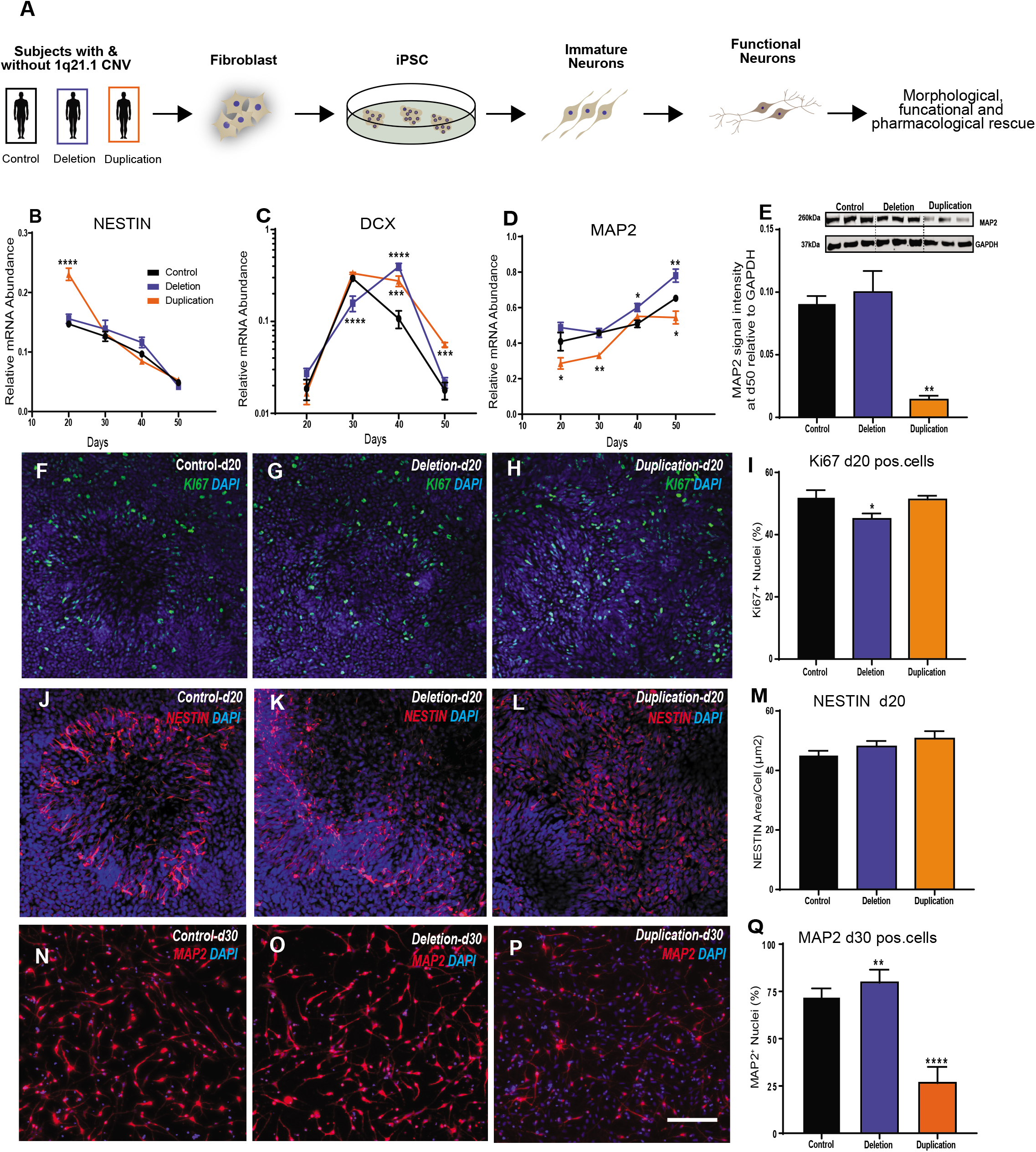
Cortical neurons with 1q21.1 deletion and duplications are associated with aberrant maturation. **A** Workflow for the generation of stable iPSC lines from human fibroblast carrying 1q21.1 deletions and duplication and the subsequent conversion of iPSC into functional neurons. **B-D** Gene expression analysis for the expression of NESTIN, DCX and MAP2 from day 20 to day 50 of neuronal differentiation in control, 1q21.1 deletion and 1q21.1 duplication neuronal culture. NESTIN, genotype (F_2,72_=7.94; P<0.001; n≥3/group), time (F_3,72_=193; P<0.0001; n≥3/group); DCX, genotype (F_2,72_=7.561; P<0.0001; n≥3/group), time (F_3,72_=120.4; P<0.0001; n≥3/group) interaction between genotype and time (F_6,72_=20.88, P<0.0001, n≥3/group); MAP2, genotype (F_2,72_=28.17; P<0.0001; n≥3/group), time (F_3,72_=53.13; P<0.0001; n≥3/group). Data sets were analysed by two-way ANOVA with post hoc comparisons using Dunnett’s multiple comparisons test comparing to control samples. Stars above points represent significance following Dunnett-corrected post hoc tests. **E** Representative western blot protein bands and quantitative analysis for MAP2 expression normalised to GAPDH (n≥3). **F-H** Example images of KI67+ staining at day 20 of neuronal differentiation in control, 1q21.1 deletion and 1q21.1 duplication cell lines. **I** Quantification of the percentage of DAPI+ nuclei which colocalized with KI67 positivity in day 20 neuronal cultures (n≥3). **J-L** Representative images of NESTIN+ cells at day 20 of neuronal differentiation in control, 1q21.1 deletion and 1q21.1 duplication cell lines. **M** Quantification of the area which staining positive for NESTIN normalised to the number of cell present in the field (n≥3). **N-P** Representative images of MAP2 positive immature neurons after 30 days of differentiation from a control, 1q21.1 deletion and 1q21.1 duplication cell line. **Q** Quantification of the percentage of DAPI+ nuclei which colocalized with MAP2 positivity in immature neuronal cultures after 30 days of differentiation (n≥3). Unless otherwise specified data was analysed using Students T-Tests. All data presented as means±SEM *P<0.05; **P<0.01; ***P<0.001 ****P<0.0001 vs. control. Scale bar = 100µm.

DCX expression in the 1q21.1 duplication group although was comparable to control during early stages of differentiation, showed an increased expression at day 40 and 50. On contrary, 1q21.1 deletion cell exhibited more complex pattern with a decreased expression compared to controls at day 30 and an increased expression compared to controls at day 40. However, the levels were comparable to controls at day 50 (Fig 1C). MAP2 expression was significantly reduced throughout neuronal differentiation of 1q21.1 duplication group compared to controls (Fig. 1D). This was further accompanied by a reduced number of MAP2+ cells at day 30 and reduction in MAP2 protein at day 50 (Fig1 Q, E). On contrary, 1q21.1 deletion experiential group demonstrated an increased MAP2 expression at both day 40 and 50 of differentiation and an increase in MAP2+ cells at day 30 (Fig. 1D, N-Q). Although we note an increase in MAP2 protein levels at day 50, it did not reach significance (Fig.1E). Neuronal cell morphologies were also examined at day 30 of neuronal differentiation when neuronal morphology first emerge^23^ (Supplementary Fig 8). Neurons with 1q21.1 deletion had smaller soma, whereas 1q21.1 duplication had an increased soma size. These results indicate that 1q21.1 CNV alter neuronal differentiation, and differences begin to emerge from day 20, the Neural Progenitor Cell (NPC) stage prior to neuronal differentiation. These results show that a complex set of gene expression and protein changes occur during neurodevelopment of 1q21.1 CNV patient cells, which are unlikely to arise from a simple acceleration or retardation of neuronal differentiation program. However, they do suggest that the 1q21.1 duplication may delay the transition from NPC to neurons, whereas a 1q21.1 deletion suppresses proliferation and promotes neuronal production.

### Neurons with 1q21.1CNVs exhibit alterations in the cortical neuronal identity

Alternation in corticogenesis has been linked to many developmental psychiatric disorders^29^ risk for which has been associated with CNVs at 1q21.1. We therefore looked at the formation of the early born deep layer neurons specifically examining the expression of CTIP2 and TBR1. The expression of both TBR1 and CTIP2 was increased in association with the deletion of the 1q21.1 region after 50 days of differentiation with similar increases in TBR1 and CTIP2 protein expression observed at day 50. These results were confirmed using immunocytochemistry and indicate an increase in the number of CTIP2+ cells in 1q21.1 deletion. Conversely, 1q21.1 duplication was associated with a transient increase of TBR1 expression at day 30 of differentiation and no significant change in CTIP2 expression. Furthermore, at the protein and cellular level, the expression of TBR1 and CTIP2 in 1q21.1 duplication cultures were comparable to controls (Fig. 2A-C, G-I). 1q21.1 deletion and duplication neurons also expressed of upper layer markers CUX1, STATB2 and REELIN as determined by mRNA expression. The level of expression was higher in neurons these neurons in compassion to controls (Supplementary Fig 7J), suggesting that 1q21.1 mutation influences cortical identity of the iPSC derived neurons.

**Fig 2.**
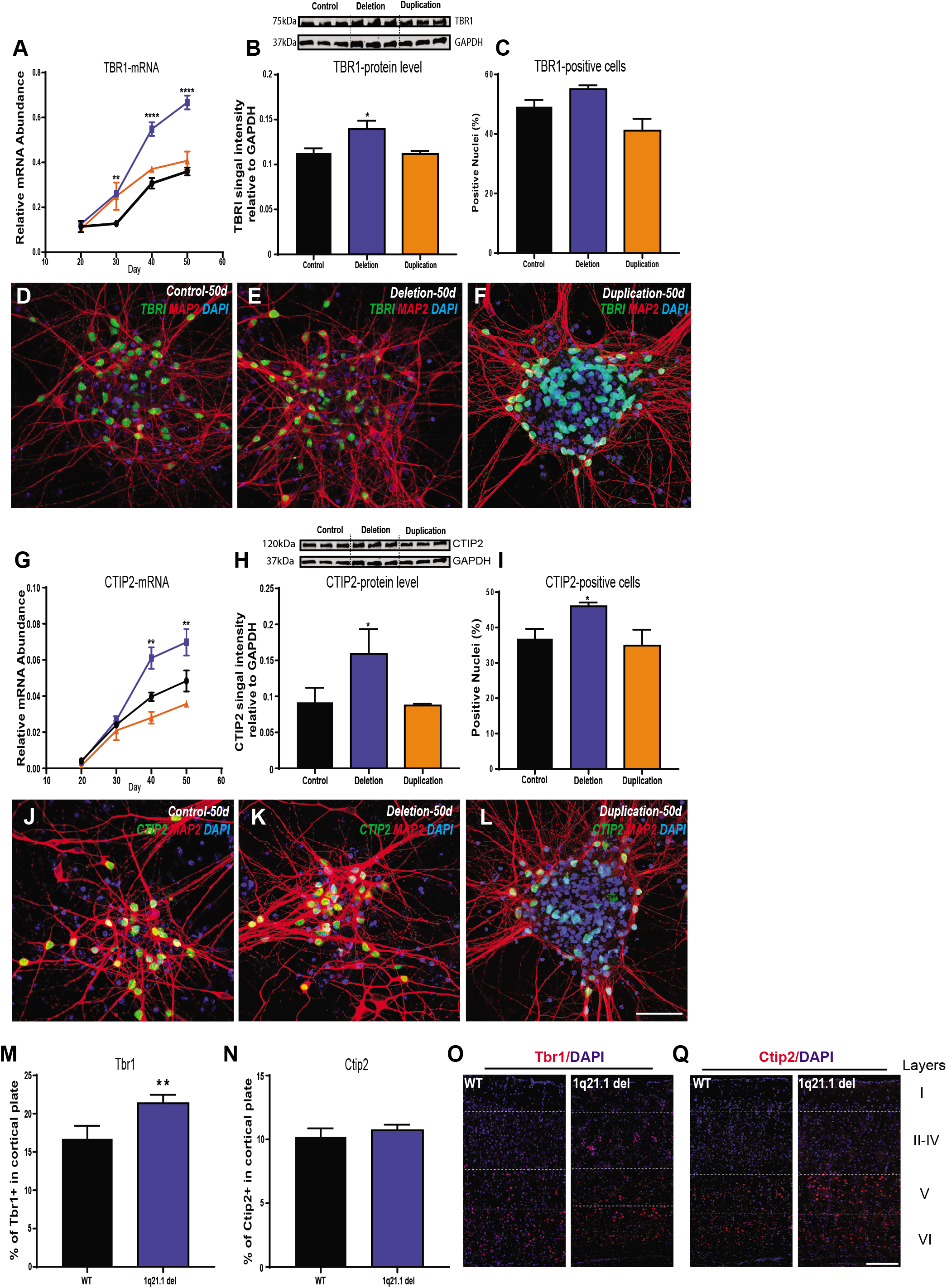
Deletion or duplication of the 1q21.1 locus is associated with aberrant cortical neuron differentiation similar to deficits found in the 1q21.1 microdeletion mouse model. **A** Gene expression of cortical deep layer marker TBR1 through the course of the neuronal differentiations (from day 20 to day 50). Both genotype (F_2,84_=57.55; P<0.0001; n≥3/group) and time (F_3,84_=365.8; P<0.0001; n≥3/group) had significant effects on TBR1 expression. **B** Representative western blot protein bands and quantitative analysis of TBR1 expression normalised to GAPDH (n≥3). **C** Quantification of the percentage of MAP2 positive cells which co-localized with TBR1 (n≥3). **D-F** Representative images of MAP2 and TBR1 colocalization from a control, deletion and duplication cell line. **G** The expression of CTIP2 through the course of neuronal differentiation (from day 20 to day 50). Both genotype (F_2,84_=199.7; P<0.0001; n≥3/group) and time (F_3,84_=133.2; P<0.0001; n≥3/group) had significant effects on CTIP2 expression. **H** Representative western blot protein bands and quantitative analysis for CTIP2 expression normalised to GAPDH (n≥3). **I** Quantification of the percentage of MAP2 positive cells which co-express CTIP2 (n≥3). **J-L** Representative images MAP2 and CTIP2 colocalization from a control, deletion and duplication cell line. Scale bar = 100µm. Data sets are expressed as mean of at least three independent experiments and were analysed by Students T-Test or two-way ANOVA with post hoc comparisons using Dunnett’s multiple comparisons test comparing to control samples. Where appropriate stars above points represent Dunnett-corrected post hoc tests. **M** Quantification of TBR1+ nuclei in the somatosensory cortex of adult mice modelling 1q21.1 deletion and wild type liter matched controls (WT) given as a percentage of nuclei in a 300µm section of cortex. **N** Quantification of CTIP2+ nuclei in the somatosensory cortex of adult mice modelling 1q21.1 microdeletion and liter matched controls given as a percentage of nuclei in a 300µm section of cortex. Data was analyzed using Student’s T-tests with 6 animals per group and n≥3 for each animal. All data presented as means±SEM *P<0.05; **P<0.01; ***P<0.001 ****P<0.0001 vs. control. **O** Representative images of a coronal brain section of wild type (WT) and 1q21.1 microdeletion model (1q21.1) showing the expression of Tbr1+ cells in the somatosensory cortex of 1-month old adult mice. **Q** Representative images of a coronal brain section of wild type (WT) and 1q21.1 microdeletion model (1q21.1) showing the expression of Ctip2+ cells in the somatosensory cortex of 1-month old adult mice. Scale Bar = 100µm.

To examine the potential effects that the changes seen in differentiating cells may have on brain organisation, we analysed 1-month old brains of a mouse model with a 1q21.1 microdeletion^30^. This analysis demonstrated that there was a significantly higher proportion of TBR1 cells in brains of the 1q21.1 mouse model in comparison to control littermates (Fig 2M). These data suggest that 1q21.1 deletion results in altered cortical patterning due to an increase in the production of lower layer cortical neurons.

### Human neurons with 1q21.1 deletion or duplication are associated with defects in synaptogenesis

Considering the altered differentiation potential associated with 1q21.1 deletion/duplication, we invesitgated the impact of the 1q21.1 CNV on synaptogenesis in our patient iPSC-derived neurons. The post-synaptic marker, PSD-95 showed a reciprocal pattern for both gene expression and protein analysis with an increased expression in 1q21.1 deletion and a decrease in 1q21.1 duplication neuronal cell (Fig 3A,B; Supplementary Fig. 9). The presynaptic marker (synaptophysin; SYN) showed an increased gene expression and number of SYN+ puncta in the 1q21.1 deletion. On the other hand, duplication of the 1q21.1 locus was associated with a decrease of SYN+ puncta and a decrease of SYN protein level (Fig. 3C) when normalised to MAP2 (eliminating differences in morphology) (Fig 3C-E). These results demonstrate that both 1q21.1 deletion and duplication are associated with defects in synapse development. It has previously shown that the presence of astrocytes influences the synapse formation in iPSC derived neurons^31^. Hence, we quantified the level of GFAP expression at day 40 and day 50, we found that level of GFAP was significantly minimal to the MAP2 expression (Supplementary Fig. 7 G, H) across groups, suggesting that altered synaptogenesis associated with 1q21.1 CNV is unlikely to be influenced by pro-maturational effect astrocytes arising from differentiation.

**Fig 3.**
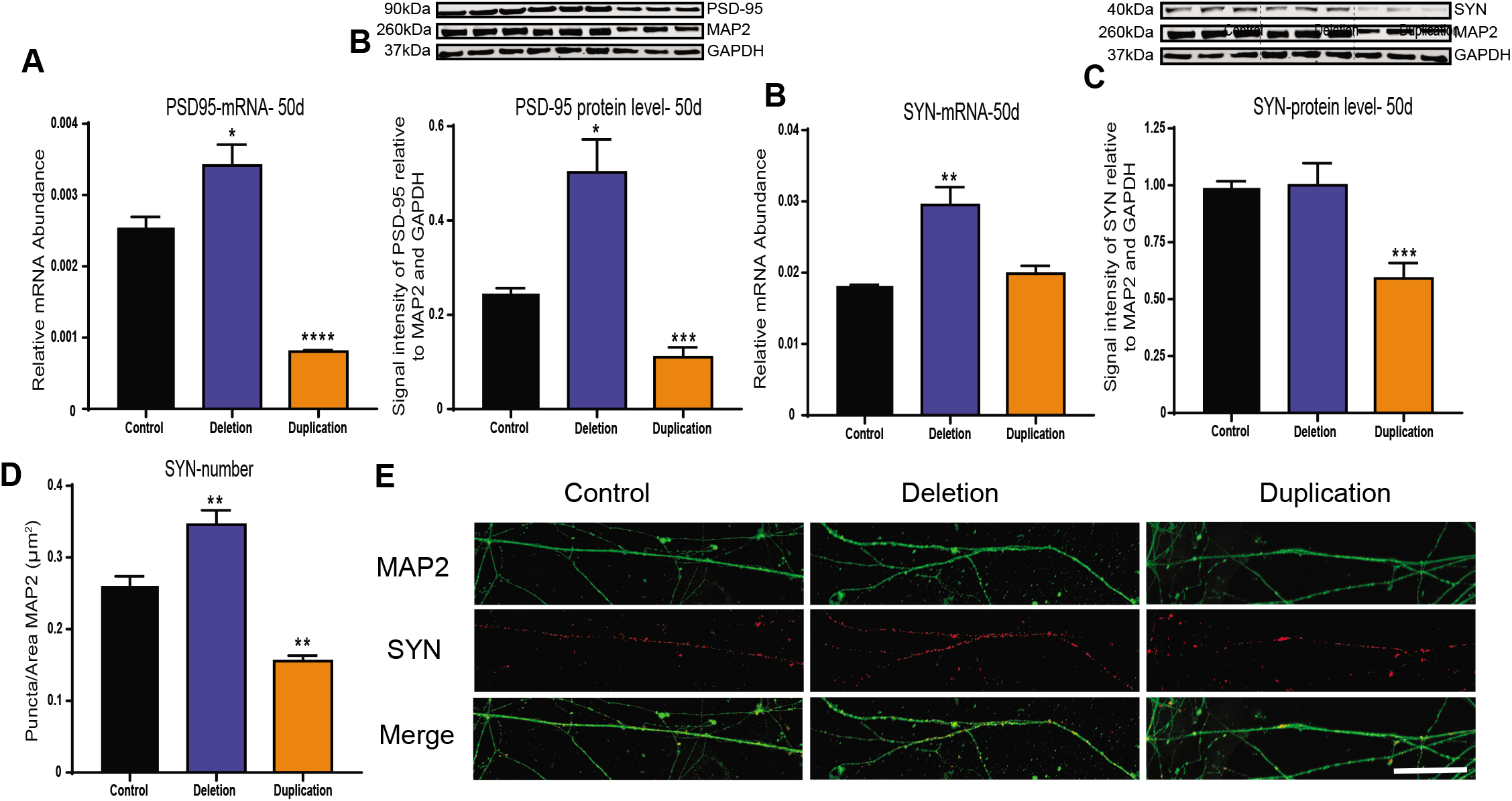
Neurons with 1q21.1 deletion and duplication are associated with synaptic defects. **A** The expression of PSD95 mRNA at day 50 of neuronal differentiation (n≥3) **B** Representative western blot protein bands and quantitative analysis for Synaptophysin expression normalized to both GAPDH and MAP2 (n≥3). **C** The expression of Synaptophysin mRNA at day 50 of neuronal differentiation (n≥3). **D** Representative western blot protein bands and quantitative analysis for Synaptophysin expression normalized to both GAPDH and MAP2 (n≥3). **E** Quantification of the number of Synaptophysin positive puncta normalised to the dendritic area stained positive for MAP2 (n≥3). **F** Representative images of Synaptophysin positive puncta in a control, deletion and duplication cell line. Scale bar = 50µm. Data was analysed using Students T-Tests and all data is presented as means±SEM **P<0.01; ***P<0.001 vs. control

### Spontaneous calcium activity reveals physiological deficits in neurons associated with 1q.21 deletion or duplications

To begin to understand the effects of 1q21.1 CNV on the physiology of neurons we assessed the cytosolic dynamics of calcium using a calcium-sensitive dye. A similar proportion of cells showed spontaneous calcium activity in the control and 1q21.1 deletion cultures (Fig. 4A). However, there were significantly fewer active neuronal cells in 1q21.1 duplication cultures compared to the controls (Fig. 4A). Quantifying the rate of spontaneous calcium activity showed a significant increase in the rate of calcium events in neurons with 1q21.1 deletion (Fig. 4B). On the other hand, after excluding the inactive cells the rate of calcium events in 1q21.1 duplication cultures was similar to the controls (Fig. 4B). Finally, the amplitude of calcium signals were comparable across the groups with no significant differences between control, deletion and duplication neurons (Fig. 4C).

**Fig 4.**
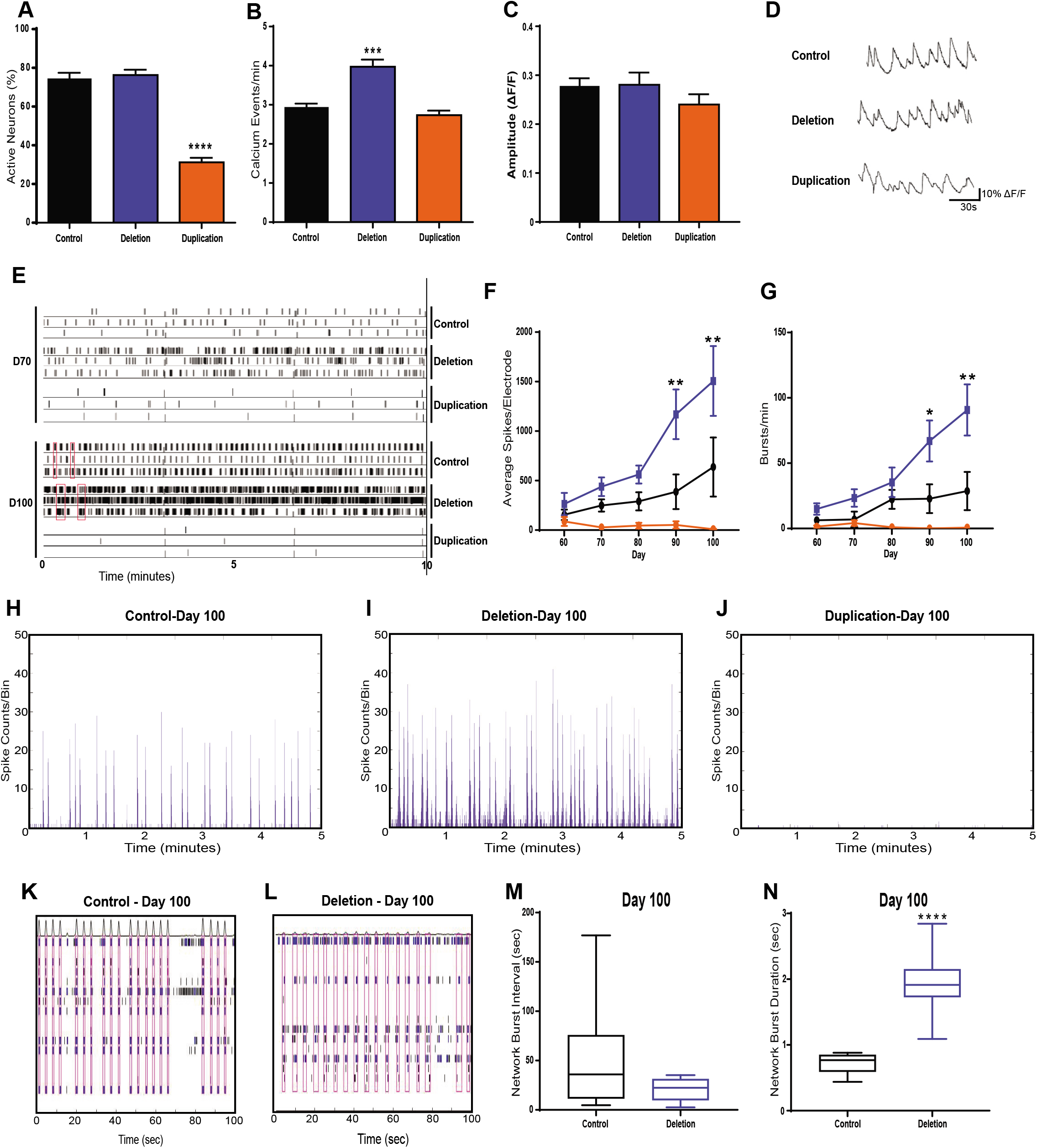
Neurons with 1q21.1 deletion and duplication display altered calcium transient activity. **A** Quantification of neuronal soma which show at least 1 characteristically neuronal calcium event (n≥3). **B** Number of characteristically neuronal calcium events recorded per minute per active neurons across different groups (n≥3). **C** Amplitude of calcium signals recorded from active neuronal cells across different groups. **D** Representative traces of calcium events as measured by changes in fluorescence from a control, deletion and duplication cell line (n≥3). **E** Representative raster plot of neuronal activity exhibited by control, deletion and duplication-derived neurons at early (D70) and late (D100) neurodevelopmental stages. Red boxes indicate periods of synchronised neuronal bursts. **F** The average number of spikes recorded per electrode across the 50 days cells were maintained on MEAs. Both genotype (F_2,76_=18.06; P<0.0001; n≥3/group) and time (F4_,76_=3.536; P<0.05; n≥3/group) had significant effects on the average number of spikes per electrode. **G** The average number bursts (defined as when more than 3 electrodes were active in the same 200ms time frame) per culture across the 50 days cells were maintained on MEAs. Only genotype (F_2,61_=8.637; P<0.001; n≥3/group) had a statically significant effects on the number of bursts per culture per recording. **H**,**I**,**J** Examples of array-wide spike detection rate (ASDR) plots, which form the basis of synchronised burst (SB) analyses. **K**,**L** Representative raster plots showing the length and interval between SBs (indicated by the red boxes) of control networks as compared to those of 1q21.1 deletion. **M** Quantification of network bursting interval in control and 1q21.1 deletion neurons (n≥3). N Quantification of network burst duration in control and 1q21.1 deletion neurons (n≥3). Data sets were analysed by Students T-Test or two-way ANOVA with post hoc comparisons using Dunnett’s multiple comparisons test comparing to control samples. Stars above points represent Dunnett-corrected post hoc tests. All data presented as means±SEM *P<0.05; **P<0.01; ***P<0.001 ****P<0.0001 vs. control

We then investigated the effect of the NMDA receptor antagonist AP5 (D-2-amino-5-phosphonopentanoate) and the AMPA receptor antagonist CNQX (6-Cyano-7-nitroquinoxaline-2,3-dione) in modulating the calcium signal in the neurons (Supplementary Fig. 10A). The addition of AP5 or CNQX resulted in a decrease in the percentage of active neurons in both the control and 1q21.1 deletion neuronal cultures. Whereas only inhibition of AMPA receptors showed a minor but significant decrease in the percentage of active neurons in 1q21.1 duplication cultures, indicating that 1q21.1 deletion neurons form connections similar to controls but the calcium activity of 1q21.1 duplication neurons could be associated with complex intricate pathways.

### Neurons with 1q21.1 deletion display neural hyperactivity, but neurons with 1q21.1 duplication display hypoactivity

The results above indicate that 1q21.1 deletion develops rapidly, expressing neurodevelopmental genes earlier than control cells, exhibiting increased synaptogenesis and increased numbers of calcium events. In contrast, 1q21.1 duplication cells showed slow or aberrant neurodevelopment, formed fewer synapses and only approx. 50% of neurons had active calcium signalling. We therefore examined the effect of the CNVs on neuronal network activity by use of Multi-Electrode Array (MEA) recordings^32^. Such networks are dependent on formation of functional synapses and are good indicators of neuronal deficits arising from aberrant neurodevelopment.

Analysis of neuronal activity over a period of 50 days post-plating onto the MEA showed that neurons with 1q21.1 deletion exhibited significantly higher spike rates and frequency of bursting compared to control neurons, particularly after D70 (Fig. 4F, G). In contrast, 1q21.1 duplication cells show no significant increases in either spike rate or burst rate during development. These data are consistent with the altered neuronal activity observed by calcium imaging. Later development time points on our MEA correspond to the emergence of large, synaptically connected neuronal networks, which burst fire in synchrony. 1q21.1 deletion patient cells exhibited synchronised bursting earlier in neuronal development (D70) than control cells (D100). Interestingly, the ultimate outcome for the neuronal network is not an increase in frequency of SBs between 1q21.1 deletion cultures and control cultures (Fig. 4M), but an increase SB duration (Fig. 4N), This aberrant network activity was inhibited by the NMDA inhibitor AP5 and the AMPA inhibitor NBQX (2,3-dihydroxy-6-nitro-7-sulfamoyl-benzo[f]quinoxaline (Supplementary Fig. 10 B-D), indicative of a glutamate transmitter dependent neuronal network. This is consistent with the higher synapse number seen in 1q21.1 deletion patient cells. In contrast, 1q21.1 duplication neurons show no neuronal network activity (Fig 4J).

### Aberrant physiological activity of neurons with 1q21.1 deletion or duplications can be rescued by modulation of Ca^2+^ activity

To determine a putative drug target to modulate the physiological deficits associated with 1q21.1 deletion and duplication we first assessed the effect of 1q21.1 mutations on the expression of neuronal ion channels. Duplication of the 1q21.1 locus was associated with a decrease in the expression of most ion channels (Fig. 5A). On the other hand, subunits of the AMPA and NMDA receptors (GLUA1 and GRIN1), and calcium channels CACNA1B and CACNA1E showed increased expression in 1q21.1 deletion neurons. To test whether a blocking calcium channel function could rescue the increased Ca^2+^ spiking in 1q21.1 deletion neurons, we added verapamil^33,34^, to neuronal cultures. Verapamil caused a significant reduction in the rate of Ca^2+^ events in 1q21.1 deletion neurons (Fig. 5C). However, there was an increased rate of Ca^2+^ events in control neurons likely caused by compensatory mechanisms due to the chronic administration of the drug. These results suggest that the blockage of calcium channels can dampen the increased rate of calcium events in 1q21.1 deletion neurons.

**Fig 5.**
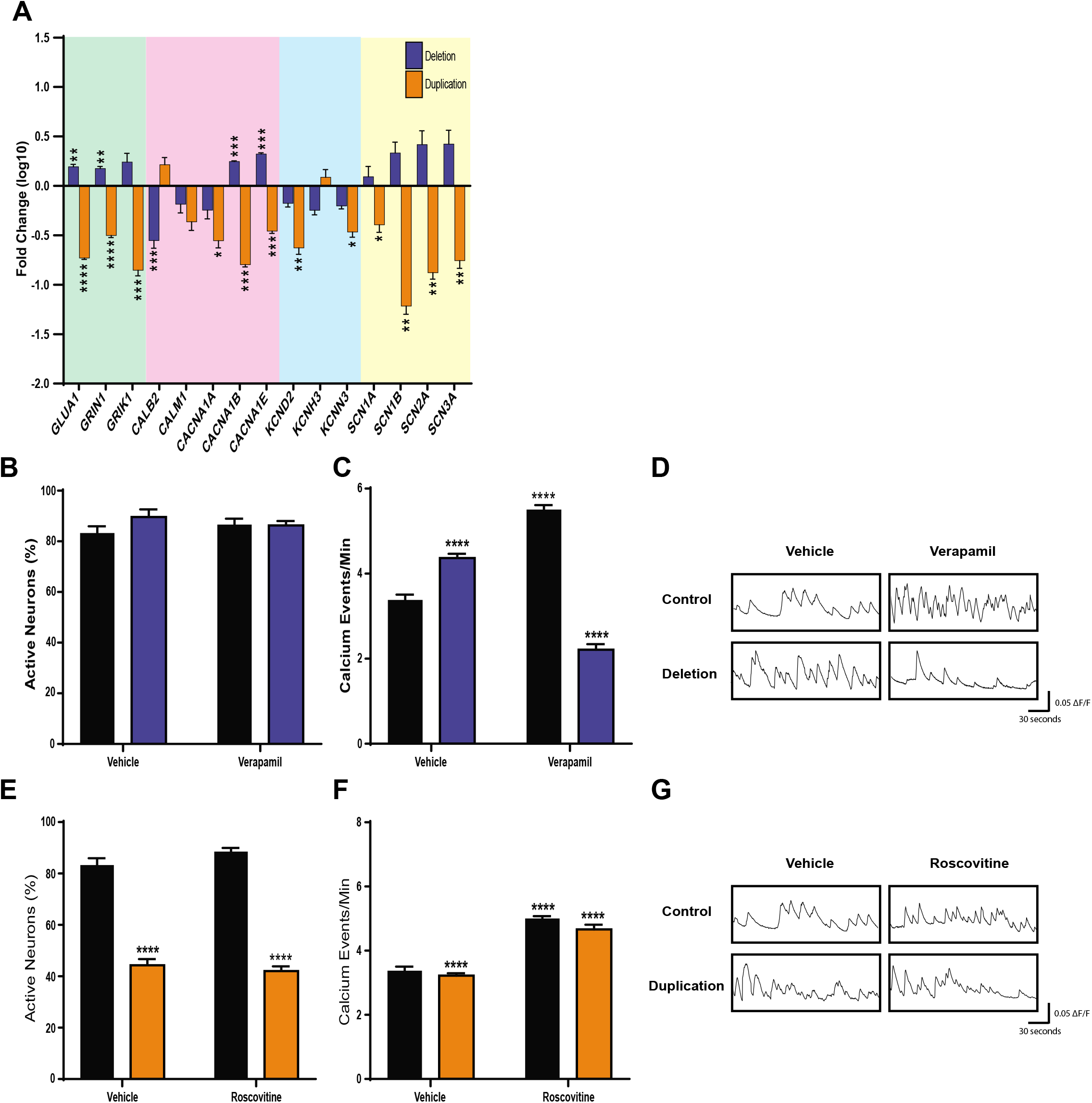
Pharmacological modulation of L-type calcium channel rescues abnormal calcium activity in neurons with 1q21.1 deletions and duplications. **A** Expression of key neuronal channels in 1q21.1 deletion and duplication neurons following 50 days of neuronal differentiation. The values are presented as fold change compared to expression in controls. Data was analysed using multiple T tests (n≥3) and significance is based on Holm-Sidak corrected P values. All data presented as means±SEM *P<0.05; **P<0.01; ***P<0.001 ****P<0.0001 vs. control. **B** Quantification of neuronal soma which show at least 1 characteristically neuronal calcium event in day 50 control and 1q21.1 deletion culture treated for 10 days with vehicle (DMSO) or verapamil (n≥3/group). **C** Number of neuronal calcium events recorded per minute in day 50 control and 1q21.1 deletion cultures treated for 10 days with vehicle (DMSO) or verapamil. Both genotype (F_1,28_=71.64; P<0.0001; n≥3/group) and the addition of verapamil (F_2,28_=79.56; P<0.0001; n≥3/group) had significant effects on the average rate of calcium events. Furthermore, there was a significant interaction between the effect of genotype and drug (F_2,28_=162.4; P<0.0001; n≥3/group) on the rate of calcium events. **D** Example traces of single neurons from both control and 1q21.1 deletion neurons treated with vehicle or verapamil. **E** Quantification of soma which show at least 1 characteristically neuronal calcium event in day 50 control and 1q21.1 deletion cultures treated for 10 days with vehicle (DMSO) or roscovitine. Only genotype had a significant effect on the percentage of active cells (F_1,22_=463.9; P<0.0001; n≥3/group). **F** Number of characteristically neuronal calcium events recorded per minute in day 50 control and 1q21.1 duplication cultures treated for 10 days with vehicle (DMSO) or roscovitine. Both genotype (F_1,22_=38.1; P<0.0001; n≥3/group) and the addition of roscovitine (F_2,22_=63.87; P<0.0001; n≥3/group) had significant effects on the average rate of calcium events. Furthermore, there was a significant interaction between the effect of genotype and drug (F_2,22_=16.06; P<0.0001; n≥3/group) on the rate of calcium events. **G** Representative traces of single neurons from both control and 1q21.1 duplication cultures treated with vehicle or roscovitine. Data sets were analysed by two-way ANOVA with post hoc comparisons using Dunnett’s multiple comparisons test comparing to control vehicle treated samples. Stars above points represent Dunnett-corrected post hoc tests. All data presented as means±SEM; ***P<0.001 ****P<0.0001 vs. vehicle treated control

To induce Ca^2+^ activity in the population of inactive cells in 1q21.1 duplication cultures, they were treated with roscovitine, which has shown to prolong the deactivation time of neuronal calcium channels^35, 36^. Addition of roscovitine significantly increased the number of calcium events in both control and 1q21.1 duplication neuronal culture (Fig. 5F). However, addition of roscovitine did not increase the proportion of spontaneously active cells in 1q21.1 duplication cultures (Fig. 5E). These results suggest that the higher and lower Ca^2+^ activity in 1q21.1 deletions and duplications can be modulated by targeting L type calcium channel antagonist and agonist.

## Discussion

One of the key findings of the study is a mirrored phenotype with respect to neuronal differentiation. Deletion of the 1q21.1 locus was associated with accelerated neuronal differentiation whereas duplication of the 1q21.1 locus had negative effects on differentiation potential. These opposing phenotypes represent a possible explanation for the micro and macrocephaly associated with CNVs at the 1q21.1 locus. In 1q21.1 deletion subjects the accelerated differentiation may result in premature loss of proliferative precursors or in premature death of new-born neurons. In 1q21.1 duplication subjects the retention of proliferative progenitors and resistance to produce mature neurons is likely result in an increase in overall cell number. However, additional work is needed in additional cellular models together with mice model to validate these explanations.

The distal 1q21.1 region consist of at least 12 protein coding genes with the recent additions of NOTCH2NLA, NOTCH2NLB and NOTCH2NLC being of particular interest^37^. A recent study investigated the effect of NOTCH2NLB on brain development demonstrated that deletion of this gene leads to premature neuronal maturation, whereas ectopic expression lead to a delay in the differentiation of radial glial cells^38^. These results are consistent with the cellular phenotypes presented in this study and provide some evidence for the underlying mechanisms involved. Given the neurodevelopment phenotype that is associated 1q21.1 CNV in our cultures it is possible that these can be attributed to dosage variation of the NOTCH2NL gene. However, the contribution of other genes within the distal 1q21.1 locus has yet to be explored; more genetic manipulation studies are needed to elucidate the contribution of each gene towards the pathology associated with 1q21.1 CNVs and it is likely that dosage level of the genes are associated stages of neuronal differentiation.

Importantly, at a functional level deletion of the 1q21.1 distal locus was associated with increase neuronal activity and deficits in neuronal network functionality (specifically in the duration of Synchronised Bursts). On the other hand, duplication of the 1q21.1 locus was associated with decreased neuronal activity and an inability to form neuronal networks. These phenotypes are in part likely a result of the altered synapse production associated with CNVs at the 1q21.1 locus. However, the cause of this synaptic disparity is not clear. Our result demonstrate that altered synaptogenesis is mediated by altered expression of synapasin. and PSD95. Future studies looking into expression analysis of these neurons will help in identifying other associated factors and in elucidating underlying common pathways (altered transcription/ altered mRNA degradation and translation), associated with synaptogenesis. Several cellular studies looking at cellular phenotypes of other CNVs such as 2p16.3/*NRXN1*, 15q13.3, 16p11.2, 22q11.21 have also shown synaptic dysfunction^39-41^. Therefore, cellular dysfunction associated with CNVs (linked to psychiatric disorders) is likely to converge on deficiencies in synaptic machinery.

Our results demonstrate that addition of verapamil (an L-type calcium channel antagonist) could reduce the rate of calcium transients in 1q21.1 deletion neurons. While some studies have questioned the specificity of verapamil targeting L-type channel^42, 43^, our results suggests that the changes in calcium movement seen in 1q21.1 deletion neurons can be modulated by altering the activity of VGCCs. The contribution and involvement of other calcium channels however cannot be ruled out. While verapamil has been used as a treatment for bipolar disorder, albeit with limited sucess^44^, it has been shown to improve scopolamine-induced memory impairments in mice^45, 46^.

Roscovitine was used in an attempt to induce calcium activity in inactive 1q21.1 duplication cells by inhibiting cell cycle progression and modulating calcium channel activity^47,35^. The addition of roscovitine was able to increase calcium activity in 1q21.1 duplication neurons consistent with previous studies^48^. However, roscovitine failed to increase the proportion of active neurons in either the control of 1q21.1 duplication group.

The present study focussed largely on identifying broad classes of neuronal dysfunction and therefore further work is necessary to elucidate the precise molecular mechanisms which underly the cellular phenotypes identified in this study. Critically future work using global transcriptomic analysis may help in identifying the precise genetic mechanisms underlying the dysfunction identified in this study.

## Funding

This work was supported by CMU fellowship to YAS. GC, AJH and JH are supported by TWF Changing Minds Programme. JH, AJH, DEJL and MO is supported by Welcome (DEFINE Strategic Award 100202/Z/12/Z). JH is also supported by Hodge Foundation (Centre Grant) and MRC grants (MR/ L010305/1, G0800509, MR/NO22572/1 and MR/L011166/1). MBMvdB is supported by MRC grants (MR/N022572/1 and MR/L011166/1). The work was also support from core facilities of the Neuroscience and Mental Health Research Institute, Cardiff University, UK and the National Center for Mental Health was supported with funds from health and Care Research Wales.

## Competing Interests

The authors declare having no competing interests

## Author contribution

Conceptualization: YAS; Planning, design and instigation: GC, YAS; Investigation, analysis and methodology: GC, MA, SL, TS, YAS; Resources including participant testing: JH, DEJL, MvdB, MO, MZ; Figures and initial writing; GC, YAS; Writing and editing of manuscript; GC, JH, AJH, YAS. All authors have read and reviewed the manuscript.

## Acknowledgements

We thank Craig Joyce and Olena Peter for their help in reprogramming cells and Dr. Faraz Mahmood Ali of the Dept. of Dermatology, University Hospital Wales for the skin biopsies.

## Notes

### Competing Interest Statement

The authors have declared no competing interest.

