## Supplemental Information for "Using induced pluripotent stem cells to investigate human neuronal phenotypes in 1q21.1 deletion and duplication syndrome"

### Supplementary Methods

#### Gene expression analysis

RNA was isolated using a GeneElute™ mammalian total RNA miniprep kit (Sigma). A minimum of 500ng of RNA was used to create cDNA using a high-capacity cDNA reverse transcription kit (Applied Biosystems). For quantitative real-time PCR 100ng of cDNA was used per reaction and primers sequences used are listed in Supplementary Table 2. Reactions were run on a StepOnePlus™ real-time PCR system (Applied Biosystems) with qPCRBIO SyGreen Blue Mix Hi-ROX (PCR Biosystems) PCR products were detected by incorporation of SYBR-green and authenticated by the melt-curves. Result were normalised to GAPDH are presented as either relative expression calculated by the $2^{-\Delta\Delta CT}$ method or by relative mRNA abundance. All samples were run in triplicate (technical replicates) and averaged before further data analysis.

#### Western Blotting

Cell lysis was performed in RIPA buffer containing protease and phosphatase inhibitors (MS-SAFE, Sigma). Protein concentration was quantified using the *DC*™ protein assay (Biorad). The samples were run on Bis-Tris 4-12% gradient gels (Thermo Fisher) and proteins were transferred to a nitrocellulose membrane. Membranes were blocked using 5% milk in tris-buffered saline (TBS) with Tween 20 (TBST). Primary antibodies (Supplementary Table 3) were diluted in blocking solution and then applied to the membranes over night at 4°C. Secondary antibodies (IRDye, Li-Cor) were applied in blocking solution for 1-1.5 hours at room temperature. Membranes were visualized using the Odessay CLx (LiCor). Quantification of bands was done using Image Studio (LiCor) and all results were normalized to GAPDH. Samples were run in triplicate (technical replicates) and were average before further data analysis.

#### Immunofluorescence and cell counting

Cells were fixed using 4% paraformaldehyde then blocked and permeabilised using 5% donkey or goat serum in PBS with 0.01-0.3% triton-X-100 (PBST). Primary antibodies (Supplementary Table 3) were applied in blocking solution and incubated overnight at 4°C. Secondary antibodies were diluted in PBST and applied for 1 hour at room temperature. Cells were counterstaining with DAPI and mounted using ProLong™ Glass Antifade Mountant (Thermo Fisher). A minimum of 3 random fields (each field containing a minimum of 20 cells) were chosen for quantification per biological replicate. Quantification was carried out using CellProfiler[22] to quantify positivity compared to DAPI. For nuclear staining, positive cells were counted only if staining overlapped with DAPI and for cytoplasmic staining positive cells were counted if staining was found in contact with DAPI. To quantify synaptic puncta Intellicount [23] was used to compare synaptic puncta to MAP2+ area using the machine learning function and an object size maxima of 9µm$^2$. All image analysis was performed on images taken using a Leica DMI600B inverted time-lapse microscope with a 20x objective.

#### Histological analysis of mice brains

Perfused and fixed brains from one month old mice modelling 1q21.1 microdeletion[24] and wild type littermates were purchased from Taconic USA (11025, n=6). Brains were incubated in 30% (wt/wt) sucrose before mounting in O.C.T (Fisher Scientific). Tissue was sliced on a cryostat at 10µm. Antigen retrieval was performed using citrate buffer. Sections were blocked and permeabilised using a solution of 5% donkey serum in PBST (with 0.1-0.5% triton-x-100). Primary antibodies (Supplementary Table 3) were applied in blocking solution overnight at 4°C. Secondary antibodies were diluted in PBST and applied for 1 hour at room temperature. Tissue was counterstained with DAPI and mounted using ProLong™ Glass Antifade Mountant (Thermo Fisher). A minimum of three sections per brain were chosen for quantification (technical replicates) and quantification was carried out using CellProfiler comparing Tbr1+ or Ctip2+ cells to DAPI in a predetermined area.

**Supplementary Fig. 1 Characterization of iPSCs generated from 1q21.1 deletion patient 1. A** Representative image of iPSCs stained for 3 markers of pluripotency (SOX2, OCT4 and NANOG). **B** Expression of OCT4 in iPSCs generated from 1q21.1 deletion patient 1 as compared to a positive control (hESCs) and a negative control (control iPSC derived neurons). **C** Expression of SOX2 in iPSCs generated from 1q21.1 deletion patient 1 as compared to a positive control (hESCs) and a negative control (control iPSC derived neurons). **D** Expression of C-MYC in iPSCs generated from 1q21.1 deletion patient 1 as compared to a positive control (hESCs) and a negative control (control iPSC derived neurons). **E** Expression of KLF4 in iPSCs generated from 1q21.1 deletion patient 1 as compared to a positive control (hESCs) and a negative control (control iPSC derived neurons). **F** Representative images and gene expression of SOX17 in iPSCs pushed to an endoderm fate. **G** Representative images and gene expression of BRACHYURY in iPSCs pushed to a mesoderm fate. **H** Representative images and gene expression of PAX6 in iPSCs pushed to an ectoderm fate. All data is presented as mean ± SEM, (n≥3) and where appropriate data was analysed by Students T-Test: ****P<0.0001 vs negative control. Scale
bar = 100µm

**Supplementary Fig. 2 Characterization of iPSCs generated from 1q21.1 deletion patient 2. A** Representative images of IPSCs stained for 3 markers of pluripotency (SOX2, OCT4 and NANOG). **B** Expression of OCT4 in iPSCs generated from 1q21.1 deletion patient 2 as compared to a positive control (hESCs) and a negative control (control iPSC derived neurons). **C** Expression of SOX2 in iPSCs generated from 1q21.1 deletion patient 2 as compared to a positive control (hESCs) and a negative control (control iPSC derived neurons). **D** Expression of C-MYC in iPSCs generated from 1q21.1 deletion patient 2 as compared to a positive control (hESCs) and a negative control (control iPSC derived neurons). **E** Expression of KLF4 in iPSCs generated from 1q21.1 deletion patient 2 as compared to a positive control (hESCs) and a negative control (control iPSC derived neurons). **F** Representative images and gene expression of SOX17 in iPSCs pushed to an endoderm fate. **G** Representative images and gene expression of BRACHYURY in iPSCs pushed to a mesoderm fate. **H** Representative images and gene expression of PAX6 in iPSCs pushed to an ectoderm fate. All data is presented as mean ± SEM, (n≥3) and where appropriate data was analysed by Students T-Test: ****P<0.0001 vs negative control. Scale
bar = 100µm

**Supplementary Fig. 3 Characterization of iPSCs generated from 1q21.1 deletion patient 3. A** Representative images of IPSCs stained for 3 markers of pluripotency (SOX2, OCT4 and NANOG). **B** Expression of OCT4 in iPSCs generated from 1q21.1 deletion patient 3 as

compared to a positive control (hESCs) and a negative control (control iPSC derived neurons). **C** Expression of SOX2 in iPSCs generated from 1q21.1 deletion patient 3 as compared to a positive control (hESCs) and a negative control (control iPSC derived neurons). **D** Expression of C-MYC in iPSCs generated from 1q21.1 deletion patient 3 as compared to a positive control (hESCs) and a negative control (control iPSC derived neurons). **E** Expression of KLF4 in iPSCs generated from 1q21.1 deletion patient 3 as compared to a positive control (hESCs) and a negative control (control iPSC derived neurons). **F** Representative images and gene expression of SOX17 in iPSCs pushed to an endoderm fate. **G** Representative images and gene expression of BRACHYURY in iPSCs pushed to a mesoderm fate. **H** Representative images and gene expression of PAX6 in iPSCs pushed to an ectoderm fate. All data is presented as mean ± SEM, (n≥3) and where appropriate data was analysed by Students T-Test: ****P<0.0001 vs negative control. Scale
bar = 100μm

**Supplementary Fig. 4 Characterization of iPSCs generated from 1q21.1 duplication patient 1. A** Representative images of IPSCs stained for 3 markers of pluripotency (SOX2, OCT4 and NANOG). **B** Expression of OCT4 in iPSCs generated from 1q21.1 duplication patient 1 as compared to a positive control (hESCs) and a negative control (control iPSC derived neurons). **C** Expression of SOX2 in iPSCs generated from 1q21.1 duplication patient 1 as compared to a positive control (hESCs) and a negative control (control iPSC derived neurons). **D** Expression of C-MYC in iPSCs generated from 1q21.1 duplication patient 1 as compared to a positive control (hESCs) and a negative control (control iPSC derived neurons). **E** Expression of KLF4 in iPSCs generated from 1q21.1 duplication patient 1 as compared to a positive control (hESCs) and a negative control (control iPSC derived neurons). **F** Representative images and gene expression of SOX17 in iPSCs pushed to an endoderm fate. **G** Representative images and gene expression of BRACHYURY in iPSCs pushed to a mesoderm fate. **H** Representative images and gene expression of PAX6 in iPSCs pushed to an ectoderm fate. All data is presented as mean ± SEM, (n≥3) and where appropriate data was analysed by Students T-Test: ****P<0.0001 vs negative control. Scale bar = 100μm

**Supplementary Fig. 5 Characterization of iPSCs generated from 1q21.1 duplication patient 2. A** Representative image of IPSCs stained for 3 markers of pluripotency (SOX2, OCT4 and NANOG). **B** Expression of OCT4 in iPSCs generated from 1q21.1 duplication patient 2 as compared to a positive control (hESCs) and a negative control (control iPSC derived neurons). **C** Expression of SOX2 in iPSCs generated from 1q21.1 duplication patient 2 as compared to a positive control (hESCs) and a negative control (control iPSC derived neurons). **D** Expression of C-MYC in iPSCs generated from 1q21.1 duplication patient 2 as compared to a positive control (hESCs) and a negative control (control iPSC derived neurons). **E** Expression of KLF4 in iPSCs generated from 1q21.1 duplication patient 2 as compared to a positive control (hESCs) and a negative control (control iPSC derived neurons). **F** Representative images and gene expression of SOX17 in iPSCs pushed to an endoderm fate. **G** Representative images and gene expression of BRACHYURY in iPSCs pushed to a mesoderm fate. **H** Representative images and gene expression of PAX6 in iPSCs pushed to an ectoderm fate. All data is presented as mean ± SEM, ($n \geq 3$) and where appropriate data was analysed by Students T-Test: ****$P<0.0001$ vs negative control. Scale bar = 100µm

**Supplementary Fig. 6 Gene expression of 1q21.1 gene**. A Schematic plot of the 1q21.1 locus. This CNV spans ~3Mb and comprises of two regions (TAR and Distal), genes known to be involved in the 1q21.1 distal/critical region (1.35Mb) are illustrated. B Bar graph showing mRNA expression changes of key genes within the 1q21.1 distal region in iPSC derived cortical neurons following 50 days of differentiation. C mRNA expression of NOTCH2NL after 20 days of neuronal differentiation. Data was analysed using Students T-Tests. All data presented as means ± SEM *$P<0.05$; **$P<0.01$ vs. control.

**Supplementary Fig. 7 Characterization of iPSC derived neurons with and without 1q21.1 CNV** . A The expression of NESTIN (NES), PLZF and ZO1 mRNA at day 20 and 50 of neuronal differentiation. The time point samples were taken had a significant effect on expression of the three markers (Control n=3, $F_{1,30}=23.2$; $P<0.0001$). The time point samples were taken had a significant effect on expression of the three markers (Control n=3, $F_{1,30}=23.2$; $P<0.0001$; ). B Fold change of NPC markers in 1q21.1 deletion and duplication cell lines after 20 days of differentiation. Data was analysed using multiple T-Tests. Stars represent Holm-Sidak corrected p-values. C, D Expression of Ki67 mRNA after 20 and 30 days of differentiation (d20 n=3; $F(2, 18) = 2.667$; $P>0.05$; d30 n=3; $F(2, 18) = 260.7$; $P>0.05$;). The expression of DCX, NCAM and MAP2 mRNA at day 20 and 50 of neuronal differentiation. The time point samples were taken had a significant effect on expression of the three markers (Control n=3; $F_{1,30}=98.4$; $P<0.0001$). Data sets were analyzed by two-way ANOVA with post hoc comparisons comparing to day 20 samples. Stars above points represent Sidak-corrected post hoc tests. F Fold change of neuronal markers in 1q21.1 deletion and duplication cell lines after 50 days of differentiation. Data was analyzed using multiple T-Tests. Stars represent Holm-Sidak corrected p-values. G The expression of GFAP was significantly minimal in comparison to MAP2 in D40 neuronal cultures. The level of GFAP was comparable across the experimental group. H Similarly, the expression of GFAP in comparison to MAP2 was significantly reduced in D50 neuronal cultures (n=3). Data was analysed using Students T-Tests. I Expression of cortical layer markers in control day 50 cultures (n=3). J Expression of cortical layer markers in 1q21.1 deletion and duplication day 50 cultures (n=3). Fold change is normalized to control day 50 differentiations. Data was analyzed using multiple t-Tests. Stars represent Holm-Sidak corrected p-values. All data presented as means ± SEM *P<0.05; ***P<0.001 ****P<0.0001 vs. control

**Supplementary Fig. 8 Morphological characterization of 1q21.1 immature neurons**. A Quantification of average soma size of neurons after 30 days of differentiation (n≥3). B Quantification of MAP2+ process length of neurons after 30 days of differentiation (n≥3). C Quantification of the number of primary MAP2+ branches of neurons after 30 days of differentiation (n≥3). Data was analysed using Students T-Tests. All data presented as means ± SEM *P<0.05; ***P<0.001 ****P<0.0001 vs. control.

**Supplementary Fig. 9 Expression of PSD-95 is reciprocally affected by 1q21.1 mutation**. A The expression of PSD-95 mRNA at day 50 of neuronal differentiation (n≥3). B Histogram of PSD-95 expression normalised to both GAPDH and MAP2 (n≥3). Data was analyzed using Students T-Tests and all data is presented as means ± SEM; *P<0.05, ***P<0.001, ****P<0.0001 vs Control. C Representative western blots of GAPDH, PSD-95 and MAP2 from a Control, Deletion and Duplication sample.

**Supplementary Fig. 10 Inhibition of AMPA or NMDA signalling results in decreased neuronal function.** A Quantification of soma which show at least 1 characteristically neuronal calcium event (n≥3). Both genotype ($F_{2,19}$=97.44; P<0.0001; n≥3/group) and drug ($F_{2,19}$=100.5; P<0.05; n≥3/group) had significant effects on the percentage of active cells. There wasalso a significant interaction between the effect of genotype and drug ($F_{4,19}$=10.16; P=0.0001; n≥3/group) on the percentage of active cells. Data sets were analysed by two-way ANOVA with post hoc comparisons using Dunnett’s multiple comparisons test comparing to control samples. Stars represent Dunnett-corrected post hoc tests. All data presented as means ± SEM *P<0.05; ****P<0.0001 vs. untreated. B Example of an array-wide spike detection rate (ASDR) plot from 1q21.1 deletion cultures after 100 days of differentiation. C Example of an array-wide spike detection rate (ASDR) plot from 1q21.1 deletion cultures after 100 days of differentiation. The culture was incubated with AP5 immediatly before recording. D Example of an array-wide spike detection rate (ASDR) plot from 1q21.1 deletion cultures after 100 days of differentiation. The culture was incubated with CNQX immediatly before recording.

**Supplementary Fig. 11** Gene expression of ion channels in control day 50 neurons. Expression of common neuronal ionchannels in day 50 control (average of both control, n≥3) neuronal culutres normalised to the expression of GAPDH.

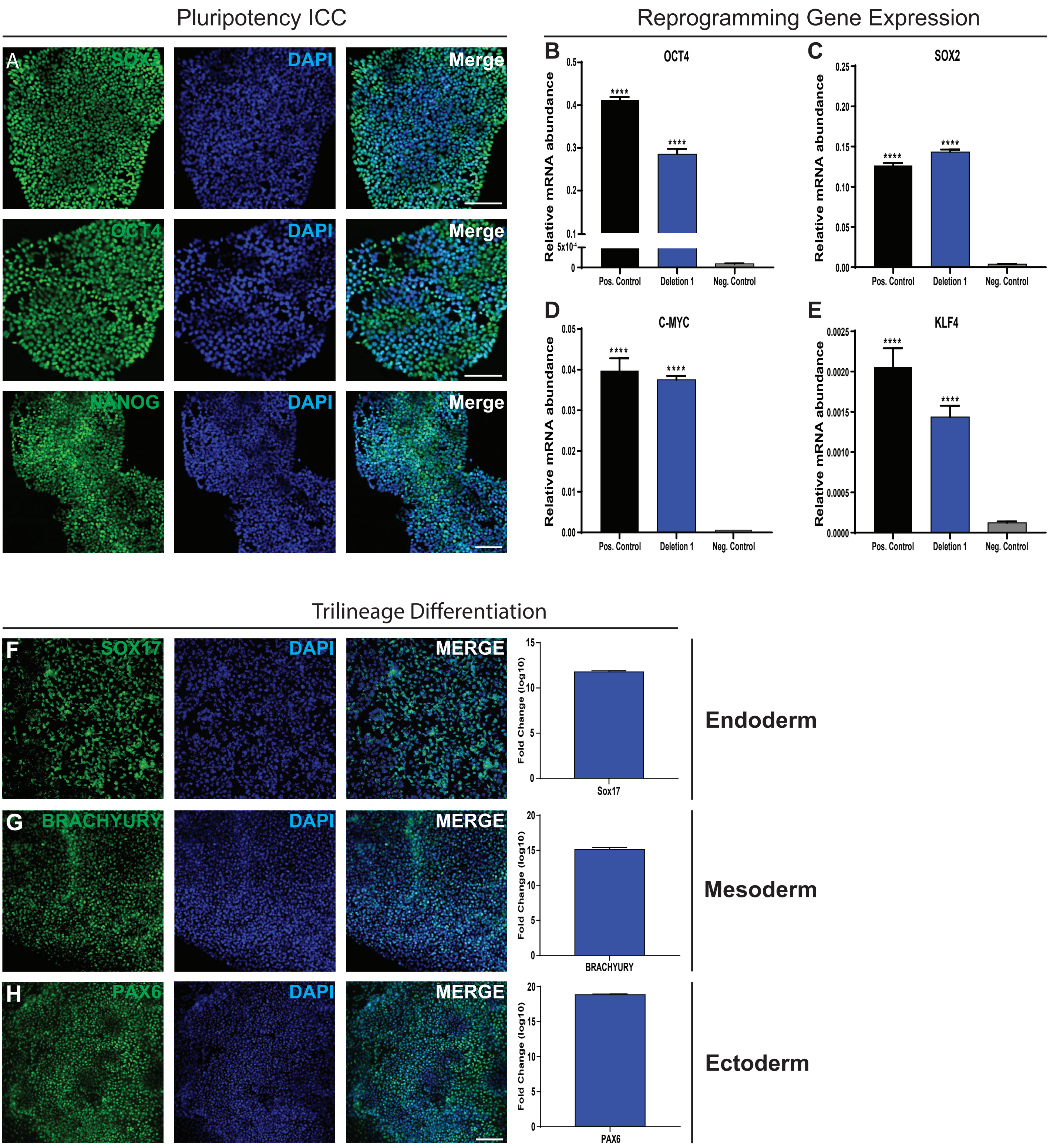


**Suppl. Fig. 1: Characterization of iPSCs generated from 1q21.1 deletion patient 1. A** Representative images of IPSCs stained for 3 markers of pluripotency (SOX2, OCT4 and NANOG). **B** Expression of OCT4 in iPSCs generated from 1q21.1 deletion patient 1 as compared to a positive control (hESCs) and a negative control (control iPSC derived neurons). **C** Expression of SOX2 in iPSCs generated from 1q21.1 deletion patient 1 as compared to a positive control (hESCs) and a negative control (control iPSC derived neurons). **D** Expression of C-MYC in iPSCs generated from 1q21.1 deletion patient 1 as compared to a positive control (hESCs) and a negative control (control iPSC derived neurons). **E** Expression of KLF4 in iPSCs generated from 1q21.1 deletion patient 1 as compared to a positive control (hESCs) and a negative control (control iPSC derived neurons). **F** Representative images and gene expression of SOX17 in iPSCs pushed to an endoderm fate. **G** Representative images and gene expression of BRACHYURY in iPSCs pushed to a mesoderm fate. **H** Representative images and gene expression of PAX6 in iPSCs pushed to an ectoderm fate. All data is presented as mean ± SEM, (n≥3) and where appropriate data was analysed by students T-Test: ****P<0.0001 vs negative control. Scale bar = 100μm.

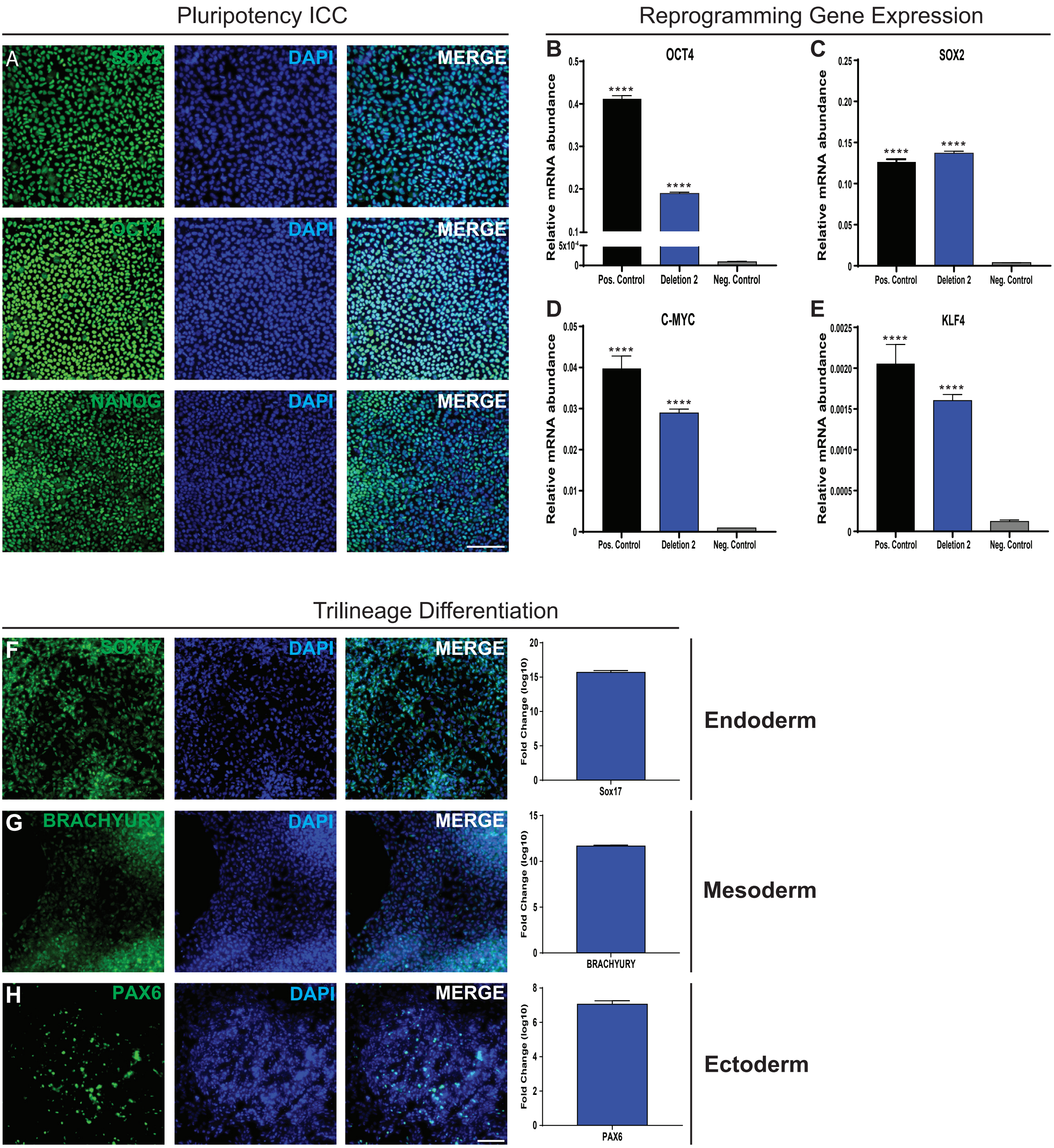


**Supp. Fig. 2: Characterization of iPSCs generated from 1q21.1 deletion patient 2. A** Representative images of IPSCs stained for 3 markers of pluripotency (SOX2, OCT4 and NANOG). **B** Expression of OCT4 in iPSCs generated from 1q21.1 deletion patient 2 as compared to a positive control (hESCs) and a negative control (control iPSC derived neurons). **C** Expression of SOX2 in iPSCs generated from 1q21.1 deletion patient 2 as compared to a positive control (hESCs) and a negative control (control iPSC derived neurons). **D** Expression of C-MYC in iPSCs generated from 1q21.1 deletion patient 2 as compared to a positive control (hESCs) and a negative control (control iPSC derived neurons). **E** Expression of KLF4 in iPSCs generated from 1q21.1 deletion patient 2 as compared to a positive control (hESCs) and a negative control (control iPSC derived neurons). **F** Representative images and gene expression of SOX17 in iPSCs pushed to an endoderm fate. **G** Representative images and gene expression of BRACHYURY in iPSCs pushed to a mesoderm fate. **H** Representative images and gene expression of PAX6 in iPSCs pushed to an ectoderm fate. All data is presented as mean ± SEM, (n≥3) and where appropriate data was analysed by students T-Test: ****P<0.0001 vs negative control. Scale bar = 100µm.

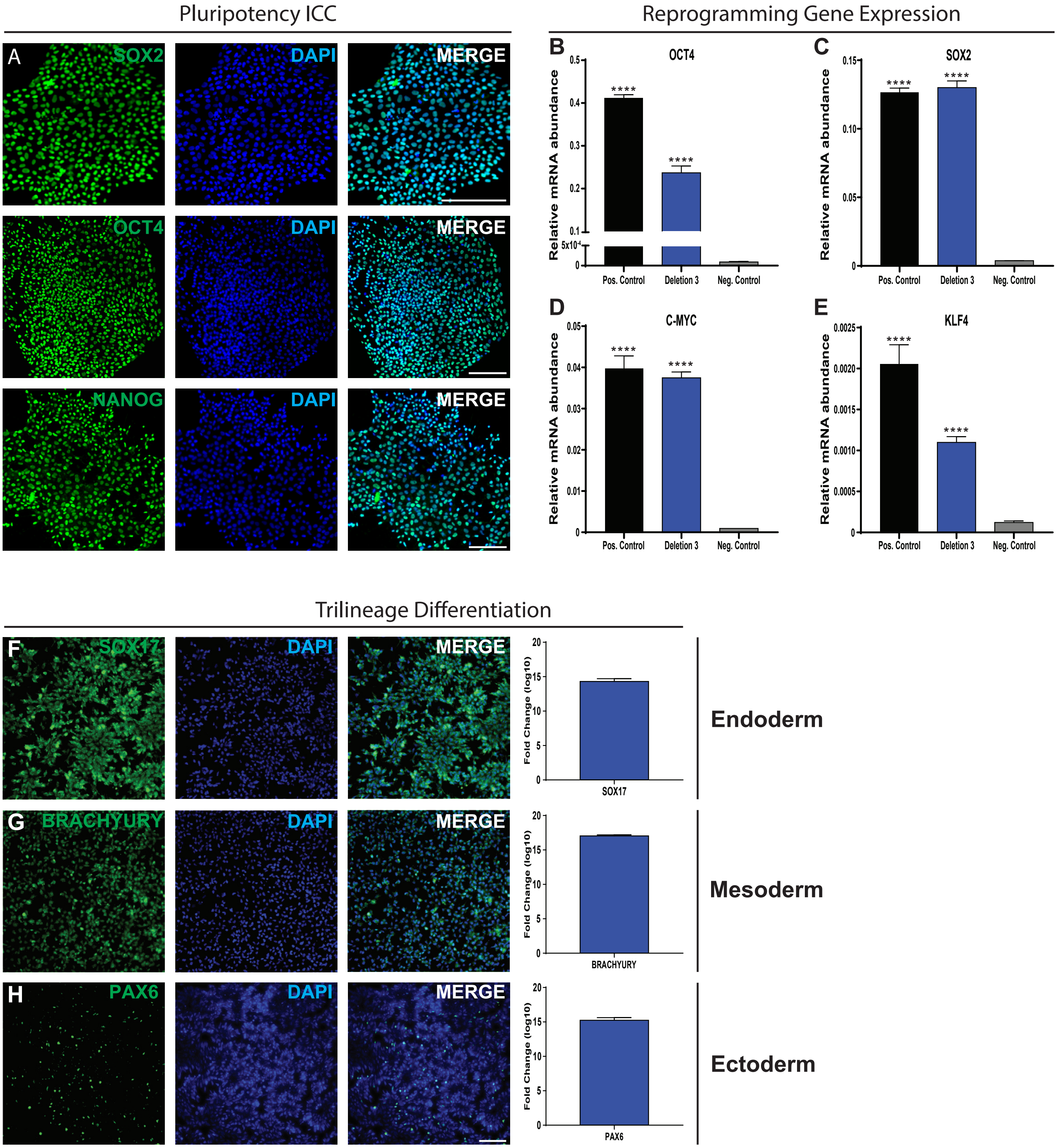


**Supp. Fig. 3: Characterization of iPSCs generated from 1q21.1 deletion patient 3. A** Representative images of IPSCs stained for 3 markers of pluripotency (SOX2, OCT4 and NANOG). **B** Expression of OCT4 in iPSCs generated from 1q21.1 deletion patient 3 as compared to a positive control (hESCs) and a negative control (control iPSC derived neurons). **C** Expression of SOX2 in iPSCs generated from 1q21.1 deletion patient 3 as compared to a positive control (hESCs) and a negative control (control iPSC derived neurons). **D** Expression of C-MYC in iPSCs generated from 1q21.1 deletion patient 3 as compared to a positive control (hESCs) and a negative control (control iPSC derived neurons). **E** Expression of KLF4 in iPSCs generated from 1q21.1 deletion patient 3 as compared to a positive control (hESCs) and a negative control (control iPSC derived neurons). **F** Representative images and gene expression of SOX17 in iPSCs pushed to an endoderm fate. **G** Representative images and gene expression of BRACHYURY in iPSCs pushed to a mesoderm fate. **H** Representative images and gene expression of PAX6 in iPSCs pushed to an ectoderm fate. All data is presented as mean ± SEM, (n≥3) and where appropriate data was analysed by students T-Test: ****P<0.0001 vs negative control. Scale bar = 100μm.

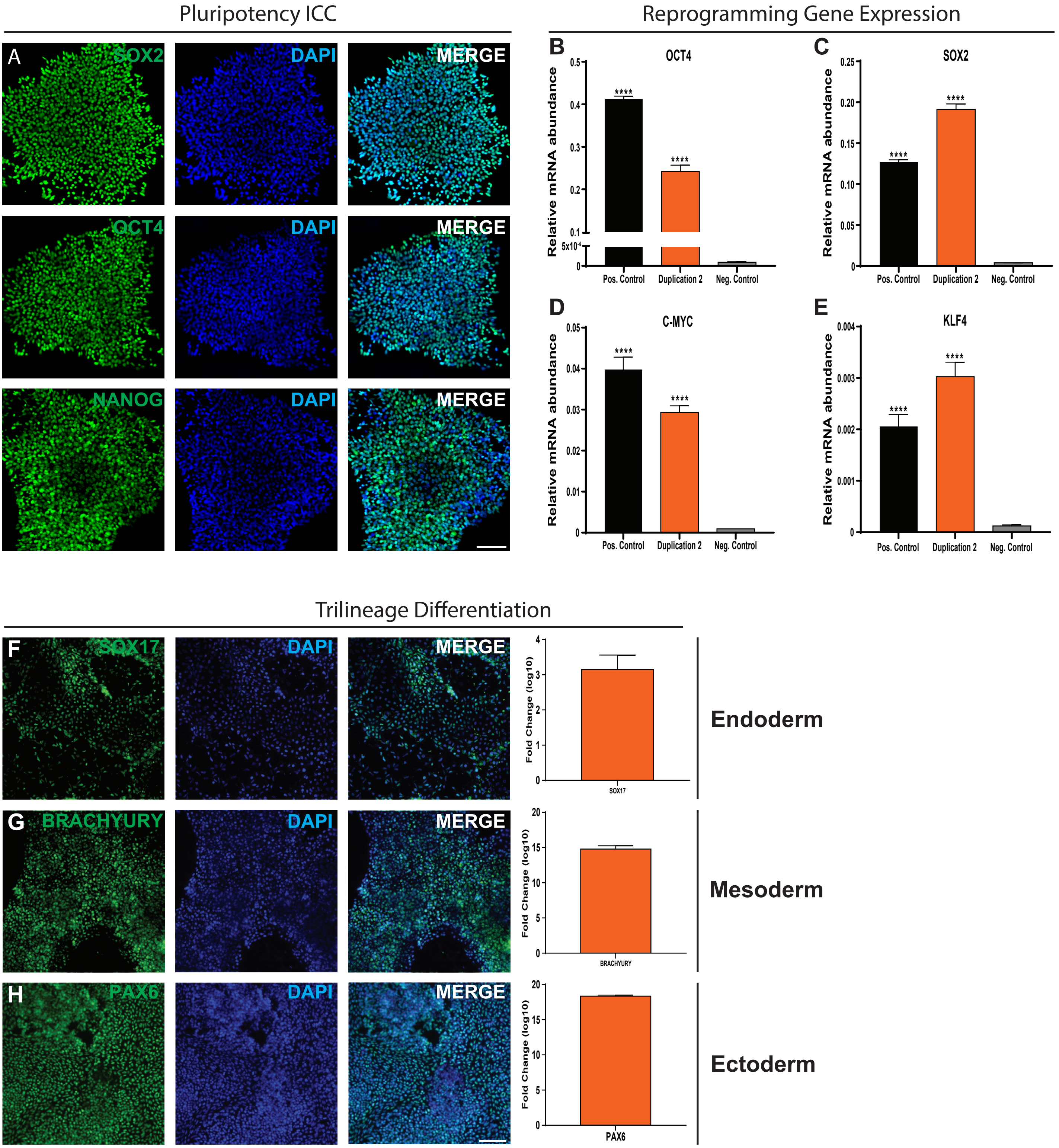


**Supp. Fig. 5: Characterization of iPSCs generated from 1q21.1 duplication patient 2. A** Representative images of IPSCs stained for 3 markers of pluripotency (SOX2, OCT4 and NANOG). **B** Expression of OCT4 in iPSCs generated from 1q21.1 duplication patient 2 as compared to a positive control (hESCs) and a negative control (control iPSC derived neurons). **C** Expression of SOX2 in iPSCs generated from 1q21.1 duplication patient 2 as compared to a positive control (hESCs) and a negative control (control iPSC derived neurons). **D** Expression of C-MYC in iPSCs generated from 1q21.1 duplication patient 2 as compared to a positive control (hESCs) and a negative control (control iPSC derived neurons). **E** Expression of KLF4 in iPSCs generated from 1q21.1 duplication patient 2 as compared to a positive control (hESCs) and a negative control (control iPSC derived neurons). **F** Representative images and gene expression of SOX17 in iPSCs pushed to an endoderm fate. **G** Representative images and gene expression of BRACHYURY in iPSCs pushed to a mesoderm fate. **H** Representative images and gene expression of PAX6 in iPSCs pushed to an ectoderm fate. All data is presented as mean ± SEM, (n≥3) and where appropriate data was analysed by students T-Test: ****P<0.0001 vs negative control. Scale bar = 100μm.

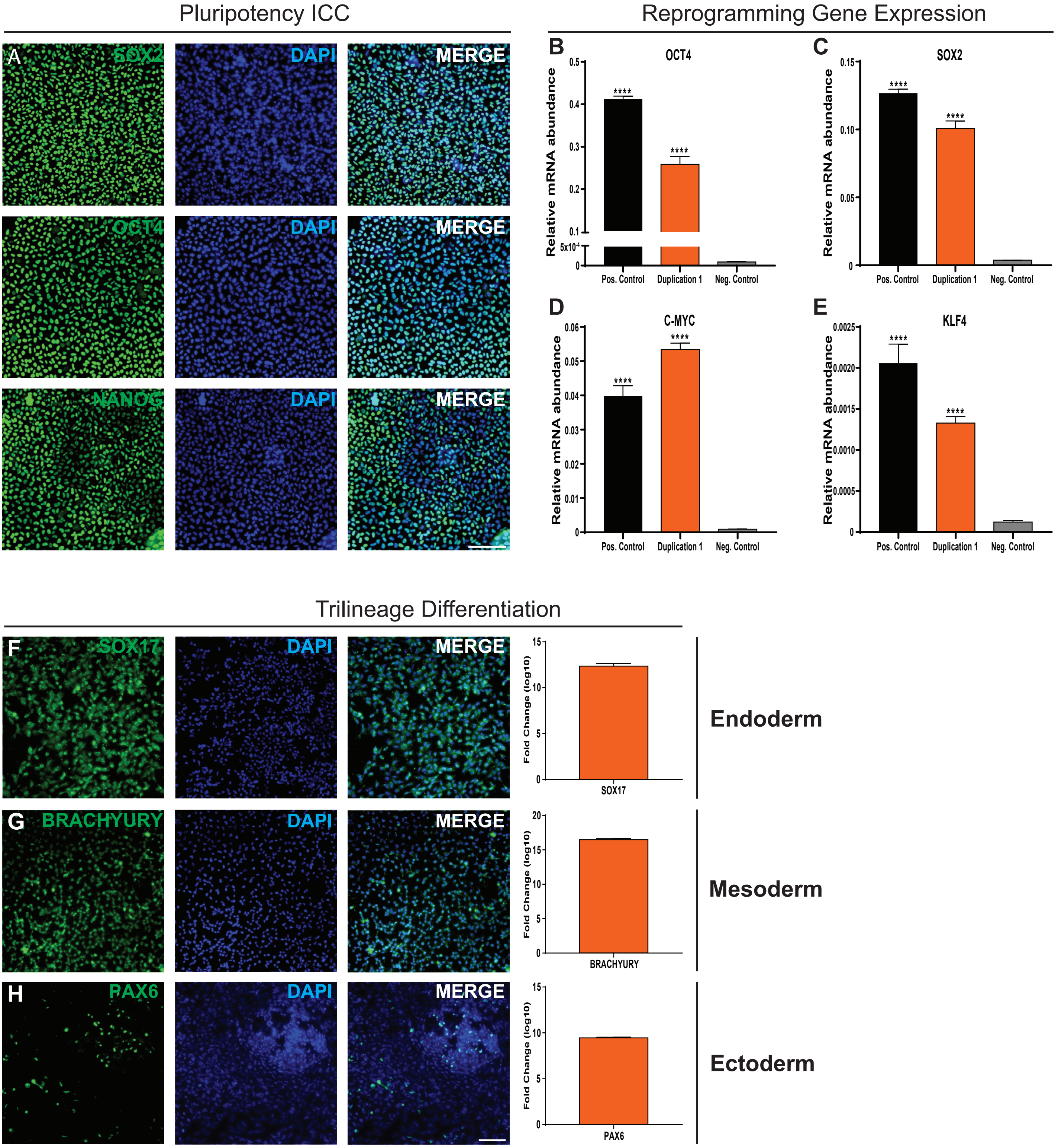


**Supp. Fig. 4: Characterization of iPSCs generated from 1q21.1 duplication patient 1. A** Representative images of IPSCs stained for 3 markers of pluripotency (SOX2, OCT4 and NANOG). **B** Expression of OCT4 in iPSCs generated from 1q21.1 duplication patient 1 as compared to a positive control (hESCs) and a negative control (control iPSC derived neurons). **C** Expression of SOX2 in iPSCs generated from 1q21.1 duplication patient 1 as compared to a positive control (hESCs) and a negative control (control iPSC derived neurons). **D** Expression of C-MYC in iPSCs generated from 1q21.1 duplication patient 1 as compared to a positive control (hESCs) and a negative control (control iPSC derived neurons). **E** Expression of KLF4 in iPSCs generated from 1q21.1 duplication patient 1 as compared to a positive control (hESCs) and a negative control (control iPSC derived neurons). **F** Representative images and gene expression of SOX17 in iPSCs pushed to an endoderm fate. **G** Representative images and gene expression of BRACHYURY in iPSCs pushed to a mesoderm fate. **H** Representative images and gene expression of PAX6 in iPSCs pushed to an ectoderm fate. All data is presented as mean ± SEM, (n≥3) and where appropriate data was analysed by students T-Test: ****P<0.0001 vs negative control. Scale bar = 100μm.

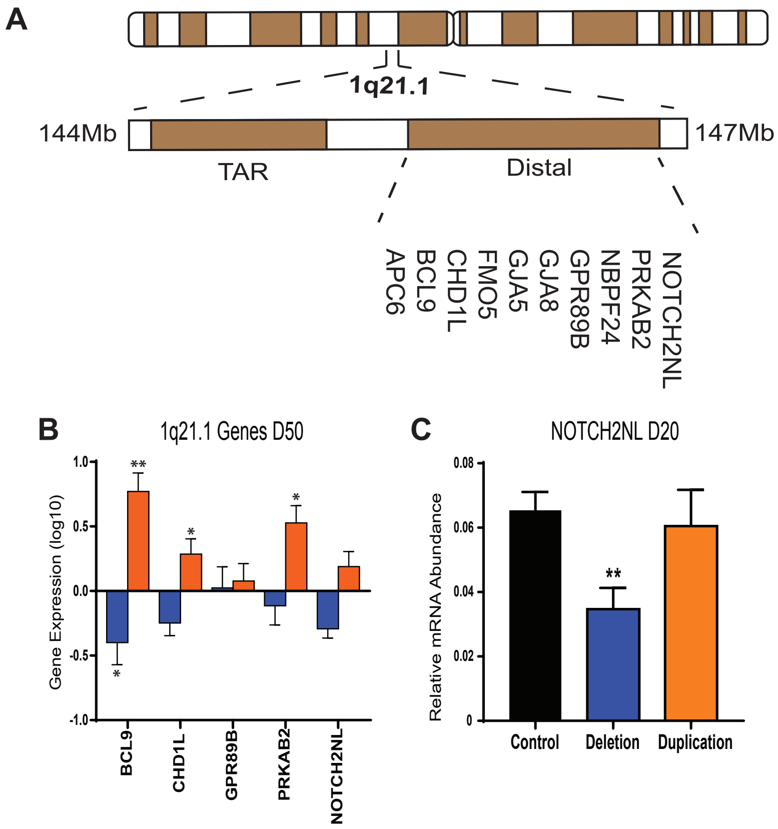


**Suppl. Fig. 6: Gene expression of 1q21.1 genes. A** Schematic plot of the 1q21.1 locus. This CNV spans ~3Mb and comprises of two regions (TAR and Distal), genes known to be involved in the 1q21.1 distal/critical region (1.35Mb) are illustrated. **B** Bar graph showing mRNA expression changes of key genes within the 1q21.1 distal region in iPSC derived cortical neurons following 50 days of differentiation. **C** mRNA expression of NOTCH2NL after 20 days of neuronal differentiation. Data was analysed using Students T-Tests. All data presented as means ± SEM *P<0.05; **P<0.01 vs. control.

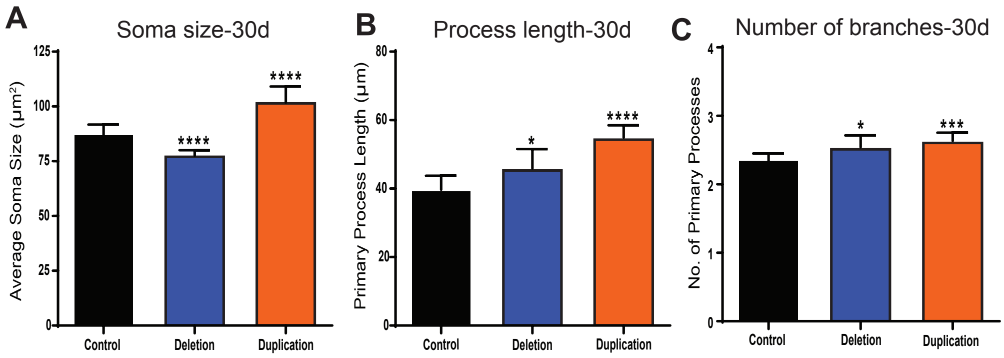


**Supp. Fig. 8: Mophological characterization of 1q21.1 immature neurons. A** Quantification of average soma size of neurons after 30 days of differentiation (n≥3). **B** Quantification of MAP2+ process length of neurons after 30 days of differentiation (n≥3). **C** Quantification of the number of primary MAP2+ branches of neurons after 30 days of differentiation (n≥3). Data was analysed using Students T-Tests. All data presented as means ± SEM *P<0.05; ***P<0.001 ****P<0.0001 vs. control.

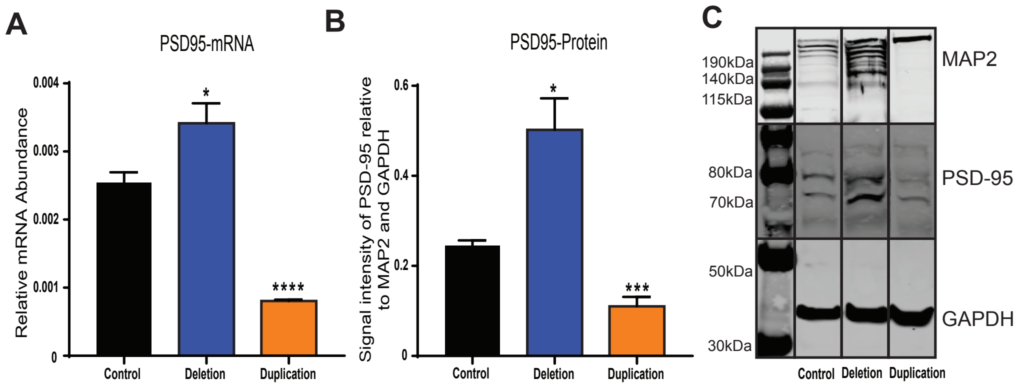


**Supp. Fig. 9: Expression of PSD-95 is reciprocally affected by 1q21.1 mutations. A** The expression of PSD-95 mRNA at day 50 of neuronal differentiation (n≥3). **B** Histogram of PSD-95 expression normalised to both GAPDH and MAP2 (n≥3). Data was analyzed using Students T-Tests and all data is presented as means ± SEM; *P<0.05, ***P<0.001, ****P<0.0001 vs Control. **C** Representative western blots of GAPDH, PSD-95 and MAP2 from a Control, Deletion and Duplication sample.

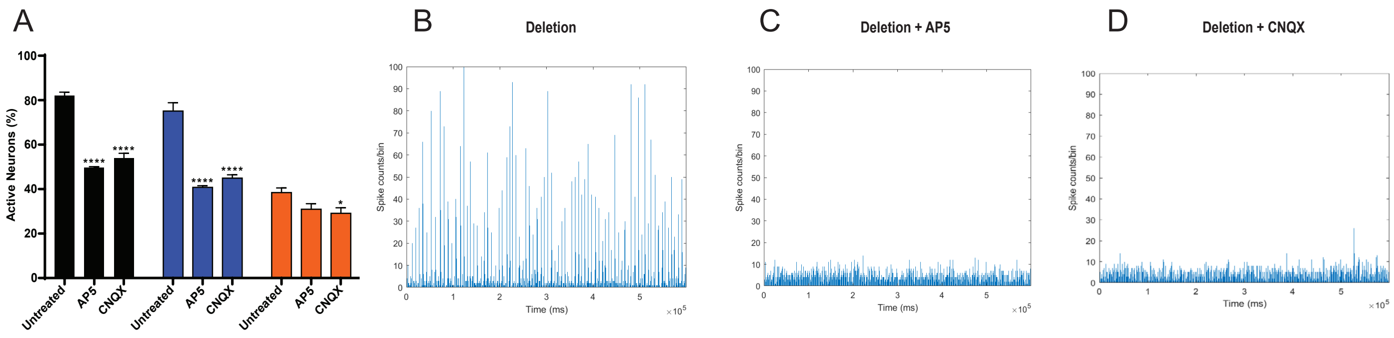


**Supp. Fig. 10:Inhibition of AMPA or NMDA signalling results in decreased neuronal function. A** Quantification of soma which show at least 1 characteristically neuronal calcium event (n≥3). Both genotype (F2,19=97.44; P<0.0001; n≥ 3/group) and drug (F2,19=100.5; P<0.05; n≥3/group) had significant effects on the percentage of active cells. There was also a significant interaction between the effect of genotype and drug (F4,19=10.16; P=0.0001; n≥3/group) on the percentage of active cells. Data sets were analysed by two-way ANOVA with post hoc comparisons using Dunnett's multiple comparisons test comparing to control samples. Stars represent Dunnett-corrected post hoc tests. All data presented as means ± SEM *P<0.05; ****P<0.0001 vs. untreated. **B** Example of an array-wide spike detection rate (ASDR) plot from 1q21.1 deletion cultures after 100 days of differentiation. **C** Example of an array-wide spike detection rate (ASDR) plot from 1q21.1 deletion cultures after 100 days of differentiation. The culture was incubated with AP5 immediatly before recording. **D** Example of an array-wide spike detection rate (ASDR) plot from 1q21.1 deletion cultures after 100 days of differentiation. The culture was incubated with CNQX immediatly before recording.

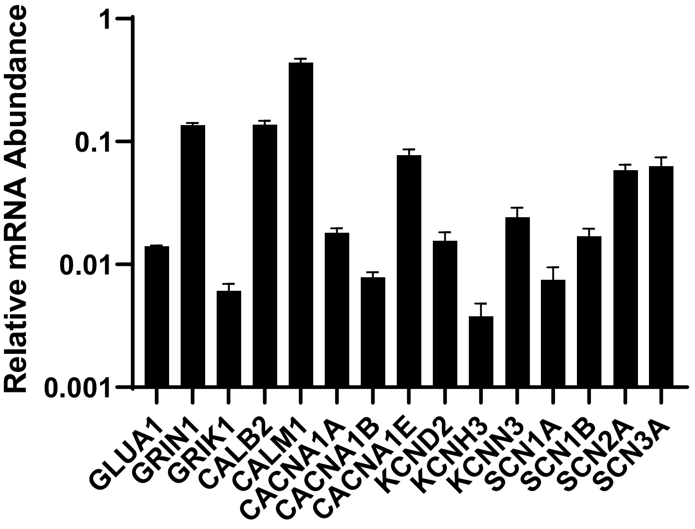


**Supp. Fig. 11:Gene expression of ion channels in control day 50 neurons.**Expression of common neuronal ion channels in day 50 control (average of both control, n≥3) neuronal culutres normalised to the expression of GAPDH.

**Supp. Table 1: Patient information for all 1q21.1 patients used in the study.** GAD: generalised anxiety disorder; MDD: major depressive; OCD: obsessive compulsive disorder; PD: personality disorder.

| | Sex | Full scale IQ | Clinical Symptoms | Medication | 1q21.1 Mutation Coordinates | Other CNVs |
|---|---|---|---|---|---|---|
| Deletion 1 | male | 77 | agoraphobia, avoidant PD, obsessive-compulsive PD, depressive PD, other substance-related disorder, conduct disorder | N/A | chr1:146496661-147375981 | chr12:2027625-2550818_dup |
| Deletion 2 | female | 78 | Adjustment Disorder, cardiac: deformed valve, narrowing of aorta, thyroid problems, hearing difficulties, sleep apnoea | Citalopram | chr1:146330584-147825662 | chr1:63918079-64038174_dup, chr5:19503096-19985234_del |
| Deletion 3 | male | 109 | social phobia, OCD, MDD, avoidant PD; interstitial cystitis (daily catheterisation), short stature, recurrent infections; tremor | Propranolol | chr1:146496661-147391614 | |
| Duplication 1 | female | 89 | social phobia, GAD, MDD, avoidant PD, conduct disorder, hearing difficulties | N/A | chr1:146330584-146982763 | |
| Duplication 2 | female | 109 | social phobia, panic disorder, agoraphobia, OCD, MDD, psychotic symptoms, schizotypal PD, Borderline PD; osteoarthritis, visual difficulties; | Propranolol | chr1:146330584-147825662 | |

**Supplementary Table 2: List of primers used in the study**

| Target | Forward | Reverse |
|---|---|---|
| BCL9 | GGCCATACCCCTAAAGCACTC | CGGAAATACTTCGCTCCCTTTT |
| CACNA1A | CGCTTCGGAGACGAGATGC | TGCGCCATTGACTGCTTGT |
| CACNA1B | GACAACGTCGTCCGCAAATAC | CCCGATGAAATAGGGCTCCG |
| CACNA1E | AATGATGCCTTAGGAGCCACC | AGCTCACGCTCAATCTGCTG |
| CALB2 | AGCGCCGAGTTTATGGAGG | TGGTTTGGGTGTATTCCTGGA |
| CALM1 | TTGACTTCCCCGAATTTTTGACT | GGAATGCCTCACGGATTTCTT |
| CHD1L | GCTATGAGCGTGTGGATGGTT | TGCTGTTAAGTTCATGCCAACTC |
| CUX1 | GCTCTCATCGGCCAATCACT | TCTATGGCCTGCTCCACGT |
| DCX | CCTTGGCTAGCAGCAACAGT | CCACTGCGGATGATGGTAA |
| ETV1 | CTGGATGACCCGGCAAATTCT | CCTCTTCAGGCTCAATCAGTTT |
| FOXP1 | AGACAAAAAGTAACGGTTCAGCC | CGCACTCTAGTAAGTGGTTGC |
| GFAP | AGGTCCATGTGGAGCTTGAC | GCCATTGCCTCATACTGCGT |
| GLUA1 | CGAGCTTTCCCGTTGATACAT | TCTGCCACTTGTAATGGTCAATG |
| GPR89B | GGAGTGACTCTCATGGCTCTT | TGTTATGCACTTCCCCCTTCT |
| GRIK1 | TCCTCTGCTATATCCTCCCTCA | CATCAGGGTTCGGTTTCTGTTA |
| GRIN1 | CTACCGCATACCCGTGCTG | GCATCATCTCAAACCACACGC |
| KCND2 | GGGTTTTTCATTGCCGTCTCT | CACAGCATACCGCTCTCCA |
| KCNH3 | TGGACGAGCACAAGGAGTTC | CGGTTCTTGGTTTCGCTGATG |
| KCNN3 | GCCTTCTCCTACACACCCTC | CTCGGGCGATCAGGTACAG |
| MAP2 | CTGCTTTACAGGGTAGCACAA | TTGAGTATGGCAAACGGTCTG |
| NESTIN | TCCAGAAACTCAAGCACCA | AAATTCTCCAGGTTCCATGC |
| NOTCH2NL | TGAGCCTTTGAAGCAGGAGG | AGATCCACATGGGGAGGGG |
| Pax6 | CAACTCCATCAGTTCCAACG | TGGATAATGGGTTCTCTCAAACTCT |
| PRKAB2 | ATGCGTTTCGATCTGAGGAAAG | GGTTCAGCATAACATGGTTGGG |
| PSD95 | AGCCCCAGGATATGAGTTGC | GATGTGTGGGTTGTCAGTGC |
| REELIN | TCCGGGACAAGAATACCATGT | CCAAATCCGAAAGCACTGGAA |
| SATB2 | CCGCACACAGGGATTATTGTC | TCCACTTCAGGCAGGTTGAG |
| SCN1A | ATGTGGAAATAGCTCTGATGCAG | AGCCCAACTGAAGGTATCAAAG |
| SYNAPTOPHYSIN | TGGTGTTCGGCTTCCTGAA | GCGGCCCAGCCTGTCT |

**Supplementary Table 3: List of antibodies used in the study.**

| Target | Supplier (code) | Host | Dilution |
|---|---|---|---|
| Oct4 | CellSignalling (C30A3) | Rabbit | 1:400 |
| Sox2 | CellSignalling (D6D9) | Rabbit | 1:400 |
| Nanog | CellSignalling (D73G4) | Rabbit | 1:200 |
| MAP2 | R and D (MAB8304) | Mouse | 1:1000 |
| TBR1 | Abcam (Ab31940) | Rabbit | 1:200 - 1:1000 |
| CTIP2 | Abcam (Ab18465) | Rat | 1:200 - 1:1000 |
| MAP2 | Abcam (Ab32454) | Rabbit | 1:1000 |
| Synaptophysin | Abcam (Ab32127) | Rabbit | 1:500 |
| GAPDH | Abcam (ab9485) | Rabbit | 1:5000 |
| PSD95 | Abcam (ab76115) | Rabbit | 1:1000 |
| Ki67 | Abcam (ab15580) | Rabbit | 1:1000 |
| Nestin | Abcam (ab105389) | Rabbit | 1:300 |
